# Structure and regulation of GSDMD pores at the plasma membrane of pyroptotic cells

**DOI:** 10.1101/2023.10.24.563742

**Authors:** Shirin Kappelhoff, Michael Holtmannspötter, Stefan L. Schaefer, Eleonora G. Margheritis, Hannah Veit, John S.H. Danial, Sebastian Strauss, Rico Franzkoch, Olympia Ekaterini Psathaki, Ralf Jungmann, Rainer Kurre, Gerhard Hummer, Jacob Piehler, Katia Cosentino

## Abstract

Gasdermin D (GSDMD) executes inflammatory cell death pyroptosis by permeabilizing the plasma membrane (PM). We introduce polymer-supported PM (PSPM) to gain access to the cytoplasmic side of the PM with imaging techniques while preserving the native PM complexity and lipid microenvironment. By combining PSPM with DNA-PAINT super-resolution microscopy we visualized, for the first time, GSDMD nanostructures directly at the PM of pyroptotic cells. We resolved diverse macromolecular architectures with ring-and arc-shaped GSDMD oligomers that enable PM permeabilization. The pyroptotically-inactive mutant GSDMD-C192A (human C191A) still interacts with the PM however fails to form pores. GSDMD expression levels affect pore density but not permeabilization ability. Finally, we identified the local PI(3,4,5)P_3_ concentration as a key regulatory element of PM permeabilization. Increase in PI(3,4,5)P_3_ levels in the PM during pyroptosis facilitates growth into large ring-shaped pores. Using molecular dynamics (MD) simulations, we identified the mechanism by which PI(3,4,5)P_3_ stabilizes the GSDMD assembly.

## Introduction

Gasdermins (GSDMs) are the final executors of pyroptosis [1, 2], a cell death program that plays a critical role in the innate immune response to pathogen attacks or endogenous insults [3-6]. GSDMs have the ability to perforate cellular membranes through cleavage-mediated activation by specific proteases [1, 2, 7, 8]. In the canonical and non-canonical pathways of pyroptosis, the GSDMD family member is activated by pro-inflammatory caspases, such as caspases-1, -4/5 [9, 10]. Cleavage by caspases allows the unleashing of the pore-forming N-terminal domain from the autoinhibitory C-terminal domain and its subsequent translocation to the plasma membrane (PM) to form pores [7, 9-16]. GSDMD pores mediate the release of inflammatory cytokines (e.g., IL -1β and IL -18) to recruit immune cells and induce cell lysis [10, 12, 13, 17]. In addition to triggering an inflammatory response and clearing infected or injured cells, excessive pyroptosis has been associated with various inflammatory and autoimmune diseases [18-20].

The cryo-EM structure of GSDMD pores in reconstituted systems suggests the assembly of a pre-pore on the membrane, which eventually inserts into the membrane to form a functional homogeneous annular pore composed of ∼33 subunits with an inner diameter of 21.5 nm [21]. In contrast, atomic force microscopy (AFM) in artificial membranes revealed arc-like, slit-like, and ring GSDMD shapes, all capable of perforating the membrane [11, 22]. This evidence suggests considerable plasticity of GSDMD pore assembly and opens the possibility of regulation by the PM context during pyroptosis to control the release of cell contents. Modulation of the density of GSDMD pores at the PM by activating membrane repair machineries to eliminate GSDMD pores [23, 24] or regulation of GSDMD functionality by posttranslational modifications [25-27] could be ways to control the release of mature IL-1β/18 through GSDMD pores while preventing cell lysis and detrimental pyroptosis. Furthermore, lipid interactions may contribute significantly to GSDMD function and membrane targeting specificity [7, 22, 28, 29]. The presence of PI(4,5)P_2_ at the PM enhances GSDMD activity and may facilitate the assembly of the final ring structures [7, 22]. PIPs may even regulate pyroptosis through a highly dynamic process of opening and closing individual GSDMD pores controlled by calcium influx and local phosphoinositide metabolism [29].

To uncover the implications of these regulatory principles in the physiological context, resolving GSDMD structures directly at the PM of pyroptotic cells and monitoring the effect of modulatory factors on pore formation is required. Superresolution fluorescence imaging techniques such as DNA point accumulation in nanoscale topography (DNA-PAINT) could provide sufficient resolution for the putative small size of GSDMD pores. However, the drastic morphological changes of the PM during pyroptosis and the highly fluorescent cytosolic background of cells expressing labeled GSDMD have made this task challenging. To tackle this challenge, we have devised polymer-supported plasma membrane (PSPM) for quantifying the structure and stoichiometry of GSDMD pores in their native PM environment with molecular resolution. PSPMs are intact, flat PM sheets generated by cells bound to polymer-coated surfaces, with the cell body subsequently removed. By combining PSPMs with DNA-PAINT and qPAINT super-resolution microscopy [30-32], we uncovered the presence of very heterogeneous GSDMD assemblies ranging from “undefined clusters” (indicative of small oligomeric assemblies) to arcs and rings that differ in size, number of subunits, and ability to permeabilize the PM. Modulation of GSDMD expression affects the overall density of GSDMD structures at the PM, but not the size or shape of the structures. Strikingly, we found a conversion of PI(4,5)P_2_ to PI(3,4,5)P_3_ at an early stage of pyroptosis and correlated this with the ability of PI(3,4,5)P_3_ to stabilize the opening of functional GSDMD ring structures. Based on atomistic molecular dynamics (MD) simulations, we propose a mechanism by which PI(3,4,5)P_3_ binding stabilizes GSDMD assembly. Taken together, our quantitative ultrastructural analysis of GSDMD pores in the PM of pyroptotic cells supports a model in which PIPs are fully entitled components of the GSDMD-mediated pyroptotic pathway and actively contribute to the regulatory complexity of the pore-formation process with major implications for the understanding of pyroptosis.

## Results

### Polymer-supported plasma membranes for the visualization of GSDMD structures in their native PM environment

Visualizing GSDMD pores directly at the PM of pyroptotic cells by advanced imaging techniques is hampered by the drastic morphological changes that cells undergo during pyroptosis, which causes cell swelling and partial detachment, and the fluorescent background of cytosolic labeled-GSDMD. To overcome these issues, we developed polymer-supported plasma membranes (PSPMs). PSPMs have the main advantage to allow the imaging of events on an immobilized and ultra-flat PM, thus allowing imaging of individual pore structures not biased by membrane curvature and partial detachment from the cell substrate. To prepare PSPMs from pyroptotic cells, HEK293T cells (lacking endogenous GSDMD) stably expressing murine GSDMD with mEGFP inserted between the N-and the C-terminal domains just before the caspase cleavage site (mGSDMD-mEGFP, as previously reported [33]), and a DmrB–Casp-1 construct for inducible activation of Casp-1 (henceforth HEK293T double stable or DS) were transfected with a HaloTag-mTagBFP-TMD transmembrane construct serving as a tethering protein (**Figure 1A**). Cells were cultured on a surface coated with poly-L-lysine-graft-polyethylene glycol functionalized with the HaloTag ligand (PLL-PEG-HTL). This ensured irreversible tethering of cell surface HaloTag-mTagBFP-TMD through the covalent HaloTag-HTL interaction while providing an ultrathin biocompatible polymer support [34]. After pyroptosis induction and formation of GSDMD pores, cells were treated for 5 min with the actin polymerization inhibitor Latrunculin B [35] and then subjected to shear forces by vigorous pipetting to remove the cell body (**Figure 1A**). For imaging by SEM, AFM and DNA-PAINT, the resulting PSPMs were fixed directly afterwards. Formation of intact PSPM was confirmed by fluorescence and scanning electron microscopy (SEM), and by AFM (**Figure 1B-D**). Strikingly, distinct, bright GSDMD-mEGFP spots associated with the PM could be discerned, supporting efficient isolation of GSDMD pores in PSPMs (**Figure 1B**). SEM and AFM confirmed a very flat topography of these PSPMs, with an average high of ∼13 nm above the polymer cushion (**Figure 1C-E**). Characteristically elevated levels at the cell edge can probably be attributed to collapsed PM from the top of the cell. However, towards the centre of the cell, PSPMs were homogeneous in their topography (**Figure 1C and D**). Next, we evaluated if changes in membrane curvature due to membrane tethering could prevent the formation of functional GSDMD structures. To this end, we compared the timing of pyroptosis in tethered *versus* untethered cells. Cells in both conditions showed comparable morphological alterations and similar timing of cell death detected by uptake of a PM impermeable dye indicating permeabilization (**Figure 1F and G**). We further tested the integrity of the PSPM by measuring membrane fluidity prior to fixation (**Extended Figure 1A-C**). For this purpose, we performed FRAP in HEK293T DS cells additionally expressing a farnesylated mCherry construct (**Extended Figure 1A**). Farnesyl-mCherry showed maximal recovery after 80 s with an average recovery half-time of 24.46±0.55 s, resulting in an estimated diffusion coefficient of 0.23±0.005 µm^2^/s (**Extended Figure 1B**). To visualize mobility across the PM sheet and to obtain a more accurate diffusion rate for membrane-associated proteins in PSPMs, we performed single-molecule tracking analysis. PSPMs allow efficient labelling of proteins facing the intracellular side. We then labelled a small portion of farnesyl-mCherry with an anti-mCherry nanobody conjugated to Dy647 to enable long-term single-molecule tracking (**Extended Figure 1C**). The labelled farnesyl-mCherry moved across the entire PM sheet with an average diffusion constant of 0.37±0.05 µm^2^/s. Taking into account intrinsic differences between cell types, this result is consistent with previous data of farnesylated EGFP in the PM of HeLa cells with a diffusion coefficient of ∼0.47± 0.46 µm^2^/s [36]. These results show that PSPMs form intact, flat, and functional PM sheets that give us direct access to the cytosolic side of the PM for imaging the native PM environment, without the requirement of harsh treatments of the cells.

**Figure 1.**
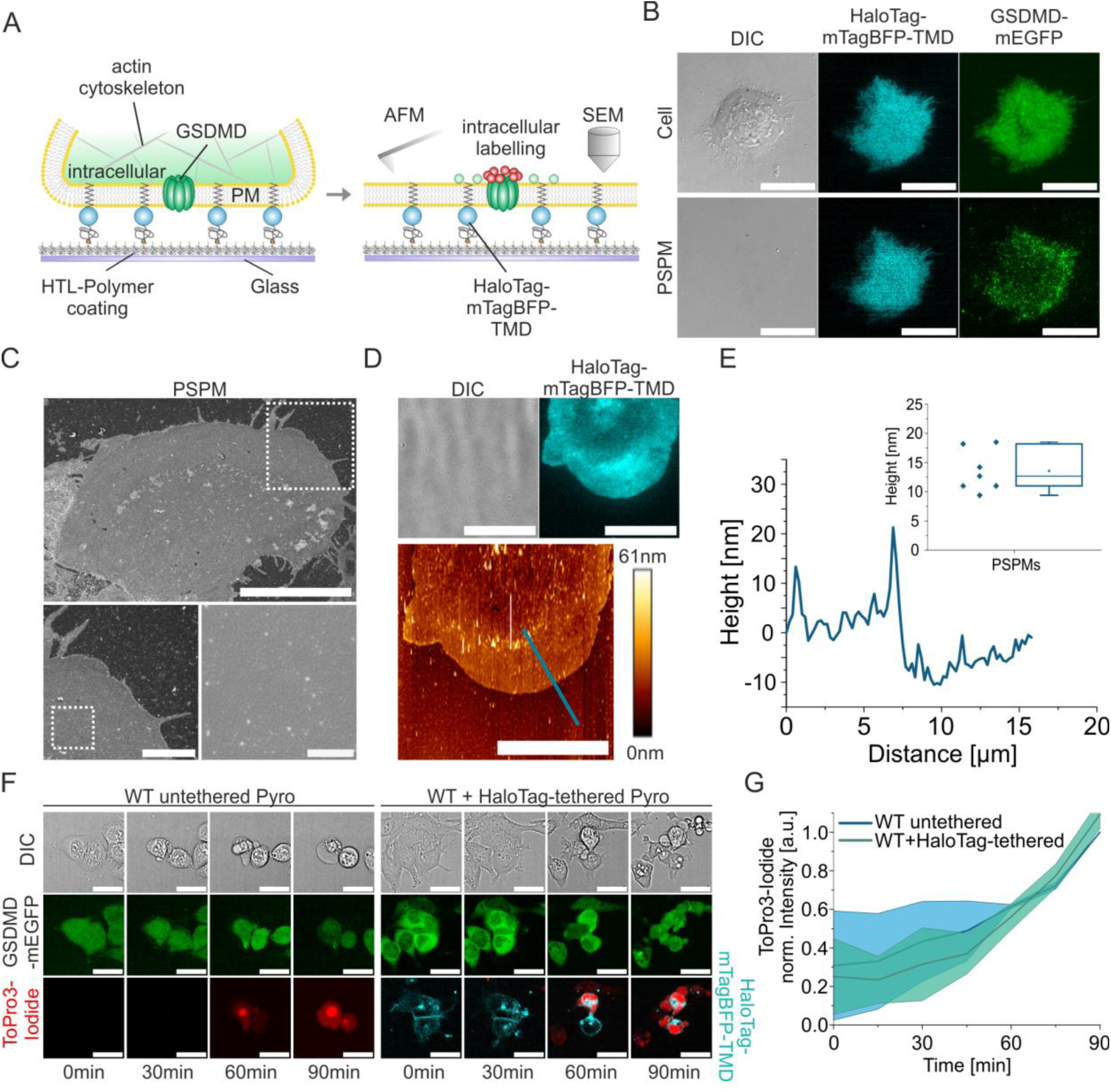
Polymer-supported plasma membranes (PSPMs) for imaging of GSDMD pores in their native PM environment. *(A) Scheme of the assay to produce GSDMD pores-containing PSPMs from pyroptotic cells for interrogation by advanced imaging techniques.* *(B) TIRF images of a representative HEK293T cell expressing the HaloTag-mTagBFP-TMD anchor (blue) and mGSDMD-mEGFP (green) before pyroptosis induction (upper row) and 90min after pyroptosis induction and PSPM generation (lower row). Scale bar 20µm.* *(C) Representative scanning electron microscopy images of a PSPM from a healthy HEK293T cell. Scale bar 20µm, 5µm and 500nm.* *(D) Representative DIC, TIRF (upper row) and AFM (lower row) images of a PSPM generated from a healthy HEK293T cell expressing the HaloTag-mTagBFP-TMD anchor protein (blue). Scale bar 20µm. The full-color height range of the AFM topograph is from low (brown-orange) to high (yellow-white).* *(E) AFM height profile of the polymer-coated surface and PSPM along the indicated line (cyan) from the AFM image in (D) and (inset) average analysis of the height of PSPMs above the polymer-coated surface (n=4 samples, 7 cells, 35 measured regions).* *(F) Representative confocal images of pyroptosis induction in HEK293T cells expressing mGSDMD-mEGFP WT without (left) and with (right) tethering by expression of the HaloTag-mTagBFP-TMD (blue) construct and cell seeding on a PLL-PEG-HTL coated surface. Pyroptosis is monitored by morphological changes (DIC) and ToPro3-Iodide staining (red). Scale bars 30µm.* *(G) Quantification of PM permeabilization of HEK293T cells as in (F) by normalized fluorescence intensity of ToPro3-Iodide over 90min (n= 2 experiments with >20 cells analyzed per experiment; Lines in the graph correspond to the average values from all measured cells and colored areas to data variability (mean ± SD)).*

### GSDMD assembles into heterogeneous nano-structures at the plasma membrane of pyroptotic cells

In addition to providing an ultra-flat membrane for unbiased imaging of pore structures, PSPMs provides direct access to the cytosolic side of the PM. We exploited this to label membrane-inserted mGSDMD-NT-mEGFP, in which the mEGFP construct faces the cytosolic PM side, with an anti-GFP nanobody carrying a DNA docking strand (GFPnb-DS) and performed DNA-PAINT super-resolution microscopy to resolve GSDMD nanostructures (**Figure 2**). For identifying pyroptotic cells after cell body removal, HEK293T DS cells expressing HaloTag-mTagBFP-TMD were cultured on a gridded and PLL-PEG-HTL-coated slide for microscopy (**Figure 2A**). Pyroptosis was induced by adding a DmrB-mCas1 dimerizer to the media [23]. After 90 min of treatment, the majority of cells underwent pyroptosis, as determined by morphological analysis and uptake of a PM impermeable dye indicating permeabilization (**Figure 2B and Extended Figure 2**). At this point, PSPMs were prepared, fixed, and incubated with GFPnb-DS for DNA-PAINT imaging (**Figure 2A, B**). Pyroptosis induction resulted in a heterogeneous signal from GSDMD-mEGFP, yet only the PSPMs clearly showed bright fluorescent dots of GSDMD at the PM in treated, but not in untreated, cells (**Figure 2B, C and Extended Figure 2B, E, F**). Super-resolution imaging of mGSDMD-mEGFP in PSPMs revealed a strong correlation of DNA-PAINT localizations with GSDMD-mEGFP signal (**Figure 2C**). After post-processing to filter out nonspecific localizations (see method section “DNA-PAINT” and **Figure 2C**), we obtained specific GSDMD signal that was not present in untreated cells (**Figure 2C and Extended Figure 2F**), corresponding to single super-resolved GSDMD nanostructures.

**Figure 2.**
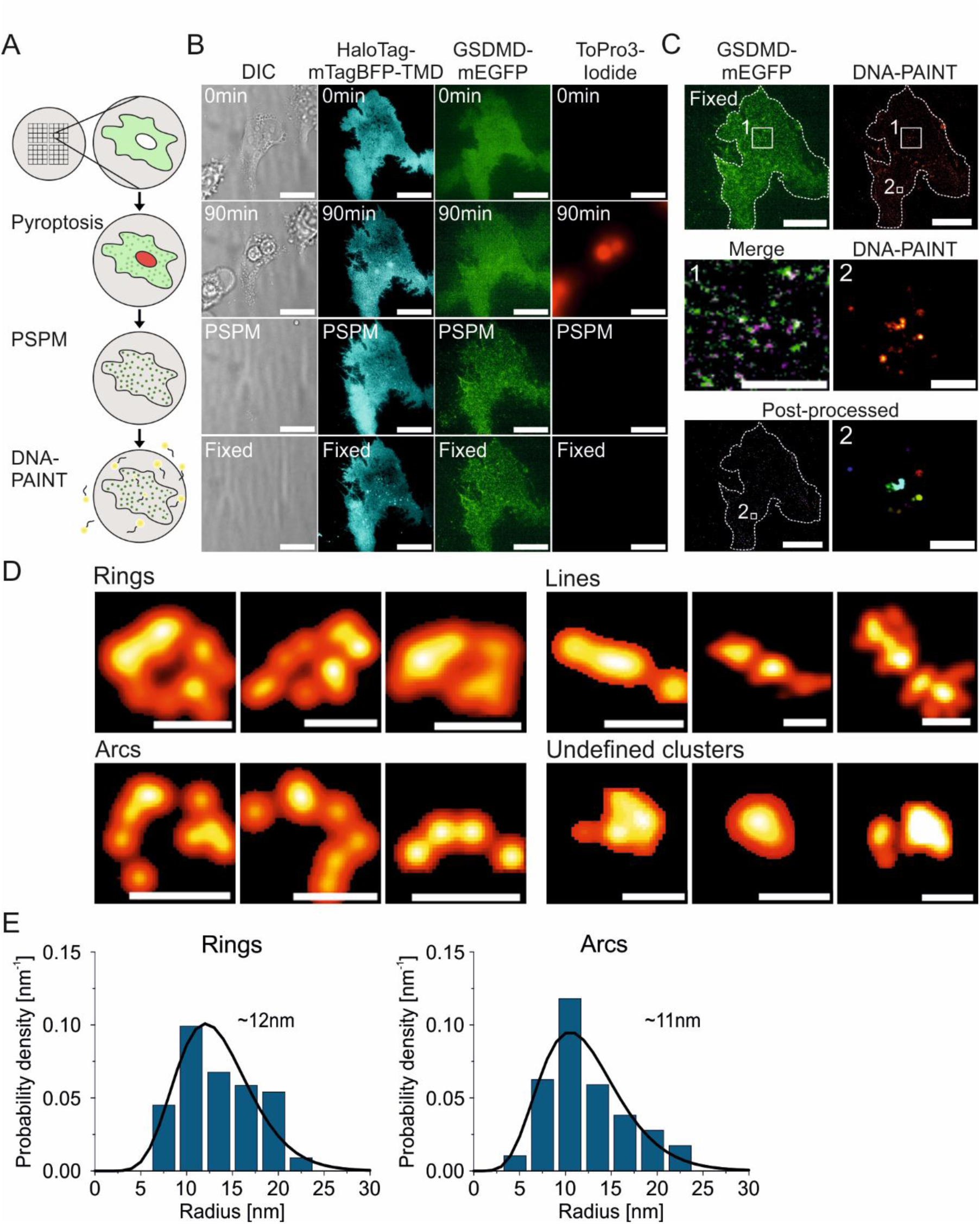
Super-resolution microscopy reveals the existence of heterogeneous GSDMD nano-structures in the PM of pyroptotic cells. *(A) Scheme of the correlative assay to monitor GSDMD structures (green) in the PM of pyroptotic cells with DNA-PAINT. HEK293T cells stably expressing DmrB-mCas1 and mGSDMD-mEGFP WT are cultured on a functionalized surface with a grid. Pyroptosis is induced by artificial dimerization of DmrB-mCas1 allowing activation of GSDMD indicated by cell permeabilization (red) and GSDMD oligomer (green dots) formation. Afterwards, PSPMs are generated allowing the removal of cytosolic fluorescent background and efficient labelling for DNA-PAINT (yellow). The grid allows identifying pyroptotic cells after cell body removal and PSPM formation for correlative DNA-PAINT imaging*. *(B) Representative TIRF images of a HEK293T cell showing the process from pyroptosis induction indicated by morphological changes (DIC), ToPro3-Iodide staining (red), and mGSDMD-mEGFP oligomers formation (green), to PSPM production (HaloTag-mTagBFP-TMD anchor, blue) and fixation for DNA-PAINT. Scale bar 20µm*. *(C) Upper row: Representative image of mGSDMD-mEGFP on PSPM (left) and corresponding DNA-PAINT image before post processing (right); Scale bars 20µm. Middle: Correlation of mGSDMD-mEGFP (green) and DNA-PAINT localizations (magenta) from zoom-in area 1 in the upper row (left, scale bar 5 µm) and zoom in of DNA-PAINT localizations from area 2 in the upper row (right, scale bar 200 nm). Lower row: post-processing image (left, scale bar 20 µm) and zoom in of area 2 (right, scale bar 200 nm)*. *(D) Gallery of GSDMD structures in PSPMs of pyroptotic HEK293T cells resolved with DNA-PAINT. Scale bars 20nm*. *(E) Quantification of the relative distribution of the radius of mGSDMD-mEGFP ring-and arc-like structures in PSPMs of pyroptotic HEK293T cells. Rings have an average radius of 12nm and arcs have an average radius of 11nm (Rings n=74 and Arcs n=102, n=8cells; 3 experiments)*.

Interestingly, we observed different macromolecular architectures rather than one type only, as would be expected if GSDMD initially arranged in a pre-pore ring structure. We classified these structures into four different shapes using the program ASAP [37]: Rings, Arcs, Lines, and Undefined Clusters, which are likely small GSDMD oligomers below the resolution limit of DNA-PAINT (**Figure 2D and Extended Figure 2G, H**).

### GSDMD ring-and arc-structures vary in size and stoichiometry

We further characterized the shape of GSDMD structures by radial profiling [37] (see method section *“Structural classification of super-resolved structures”*) and found that arcs and rings had a surprisingly broad size distribution (**Figure 2E**). Ring-like GSDMD structures exhibited a radius in the range of 7-22 nm, with an average radius of 12 nm. Arcs showed a similar distribution with an average radius of 11 nm (**Figure 2E**). Considering the size of the fused mEGFP and attached nanobody, these values are consistent with the size of GSDMD arcs and rings in reconstituted systems reported previously [11, 21, 22]. To determine the number of subunits composing GSDMD structures, we applied qPAINT, a method based on the frequency of DNA-PAINT localizations [31]. This method requires the use of DNA origami coupled to the same docking strand as the anti-GFP nanobodies as a calibration tool. For more precise quantification, the DNA origamis were deposited on the same substrate used for preparing PSPMs. For this purpose, PLL-PEG-biotin was included into the PLL-PEG-HTL surface coating to enable capturing of DNA origamis via streptavidin (**Figure 3A-C**). This approach allowed to simultaneously image DNA origamis and GSDMD structures in PSPMs by DNA-PAINT (**Figure 3C**). Comparing the binding frequency of each GSDMD structure with the average binding frequency of the monomeric binding sites of the origami structures in the same experiment, we found that the ring-shaped GSDMD structures exhibited a great diversity in stoichiometry, ranging from 8-35 subunits with an average number of 18 subunits **(Figure 3D**). The arc-shaped GSDMD structures exhibited the same heterogeneity, but with lower stoichiometry (ranging from 5-25 subunits with an average number of 14 proteins; **Figure 3D**). Overall, these results demonstrate that GSDMD is capable of forming highly heterogeneous structures that differ not only in shape but also in size. This supports the hypothesis of pores that can be appropriately modulated for selective content release.

**Figure 3.**
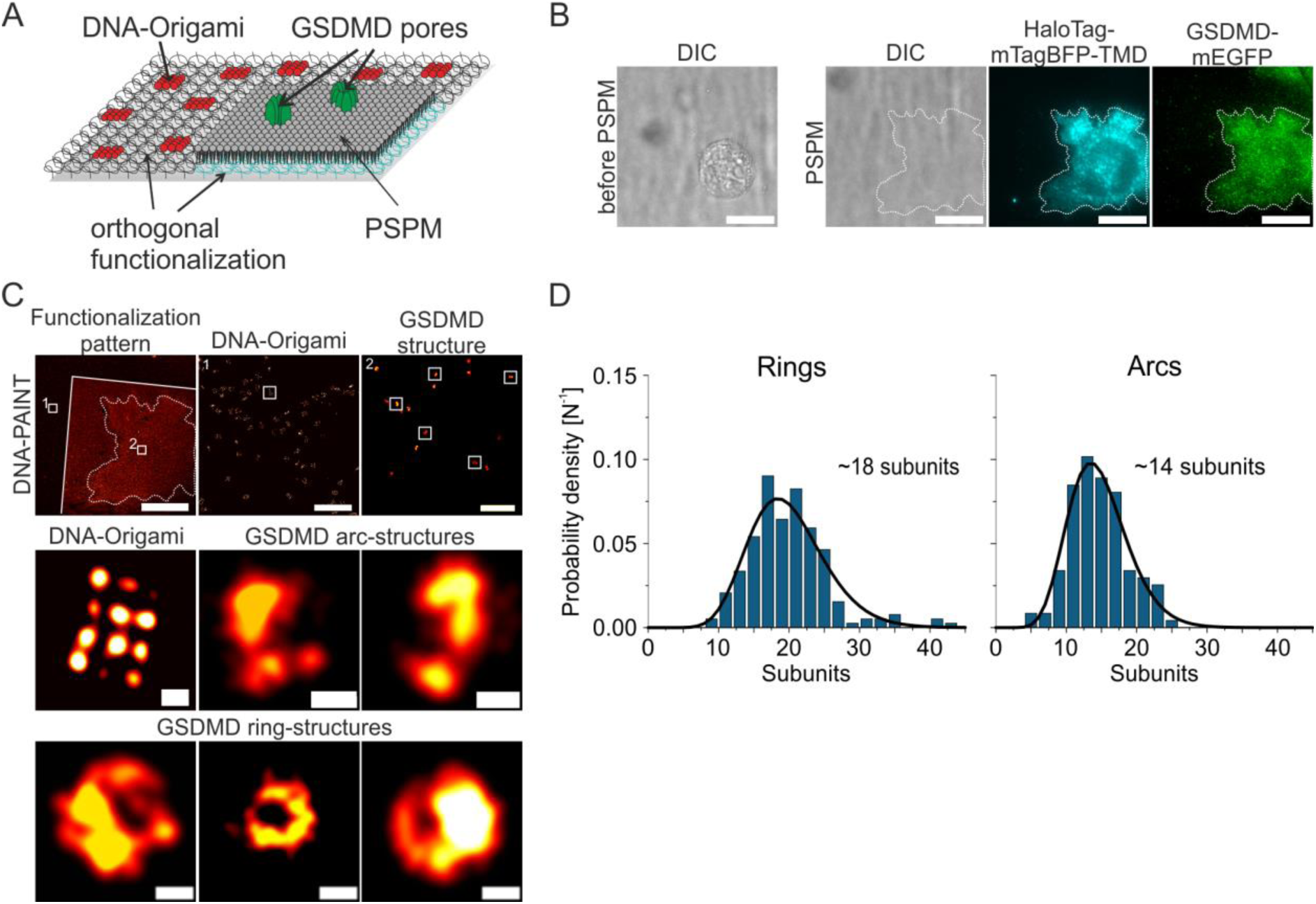
GSDMD ring and arc structures in the PM show a broad stoichiometry distribution. *(A) Scheme of orthogonal surface functionalization for simultaneous DNA-Origami immobilization (red) and tethering of PSPMs of pyroptotic cells for stoichiometry analysis of super-resolved GSDMD structures (green) by qPAINT*. *(B) Representative TIRF images of a HEK293T cell grown on an orthogonal functionalized surface before (left) and after (right) PSPM generation (HaloTag-mTagBFP-TMD anchor, blue) with mGSDMD-mEGFP oligomers (green). Scale bar 20µm*. *(C) Upper row: DNA-PAINT localizations of the PSPM shown in (B) together with immobilized DNA-Origamis serving as a calibration for qPAINT. Overview of all DNA-PAINT localizations of the patterned surface (left, scale bar 20µm) and zoomed in images of DNA-Origami (1, middle) and GSDMD structures (2, right) after post-processing (Scale bar 500nm). Middle and lower row: Representative super-resolution images of post-processed DNA-Origami and GSDMD ring-and arc-structures, (scale bar 20nm)*. *(D) Quantification of the relative distribution of the number of mGSDMD-mEGFP subunits in ring-and arc-like structures in PSPMs of pyroptotic HEK293T cells analyzed by qPAINT. Rings have on average 18 subunits and arcs have on average 14 subunits. (Rings n=194 and Arcs n=118, n=8cells; 3 experiments)*.

### GSDMD ring-and arc-structures are responsible for PM permeabilization in pyroptosis

The finding that GSDMD oligomers exist in different sizes and shapes at the PM raised the question which structures are relevant for cell permeabilization during pyroptosis. To answer this question, we compared mGSDMD-mEGFP WT with the inactive mutant mGSDMD-C192A-mEGFP at a similar expression level **(Extended Figure 3A)** [38]. Confocal microscopy of mGSDMD-C192A-mEGFP HEK293T cells after pyroptosis induction showed marked impairment of cell death (**Figure 4A, B**). Surprisingly, PSPMs prepared from cells expressing mGSDMD-C192A-mEGFP showed GSDMD dots, indicating that this mutant is still able to bind to the PM (**Figure 4C and Extended Figure 3B**). Indeed, the density of mGSDMD-C192A-mEGFP assemblies at the PM was comparable to the WT (**Figure 4D**). However, analysis of the nanoscopic structures formed by the mutant revealed striking differences from the WT, with a marked decrease in the formation of arcs (∼3% for C192A *vs*. ∼37% for WT) and ring structures (∼3% for C192A *vs.* ∼28% for WT) and a clear dominance of undefined clusters (∼87% for C192A *vs*. ∼16% for WT), which we interpret as small oligomers beyond the resolution limit of DNA-PAINT (**Figure 4E**). Overall, these data suggest that arcs and rings are the functional structures in pyroptosis. Remarkably, cells expressing mGSDMD-C192A still showed mild permeabilization (**Extended Figure 3C**), possibly due to the low proportion of arcs and rings. Whether small GSDMD oligomers are able to permeabilize the membrane for the passage of ions, as previously suggested [28, 29, 39], remains to be determined.

**Figure 4.**
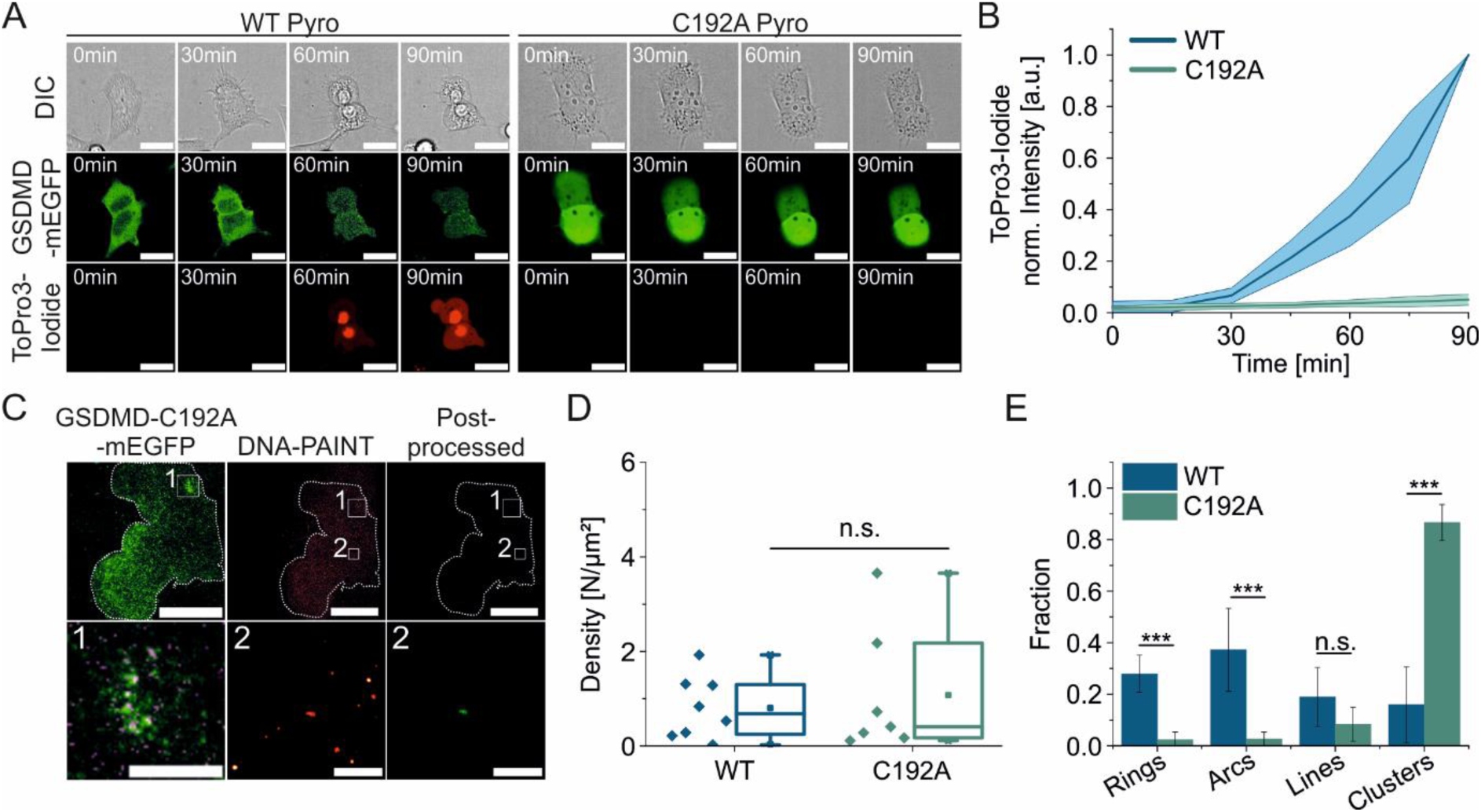
Formation of GSDMD ring- and arc-structures is associated with PM permeabilization. *(A) Representative confocal images of pyroptosis induction in HEK293T cells expressing mGSDMD-mEGFP WT (left) or the inactive mutant mGSDMD -C192A-mEGFP (right). Pyroptosis is monitored by morphological changes (DIC) and ToPro3-Iodide staining (red). Scale bars 20µm*. *(B) Quantification of PM permeabilization of HEK293T cells as in (A) by normalized fluorescence intensity of ToPro3-Iodide over 90min (n= 3 experiments with >20 cells analyzed per experiment; Lines in the graph correspond to the average values from all measured cells and colored areas to data variability (mean ± SD))*. *(C) Upper row: overview of mGSDMD-C192A-mEGFP (left) and DNA-PAINT localizations before (middle) and after post-processing (right) (Scale bars 20µm). Lower row: Correlation of mGSDMD-C192A-mEGFP (green) and DNA-PAINT localizations (magenta) from zoom-in area 1 in the upper row (left, scale bar 5 µm), zoom in of DNA-PAINT localizations from area 2 in the upper row before (middle) and after (right) post-processing. Scale bar 200 nm*. *(D) Density of super-resolved GSDMD structures after post-processing of DNA-PAINT localizations in HEK293T cells expressing mGSDMD-mEGFP WT or GSDMD-C192A-mEGFP (GSDMD WT n=8 cells; 3 experiments, GSDMD-C192A n=7 cells; 3 experiments, p=0.617)*. *(E) Comparison between the relative distribution of the different GSDMD structure types in HEK293T cells expressing mGSDMD-mEGFP WT (n=8 cells, 3 experiments, 493 structures) or mGSDMD-C192A-mEGFP (n=7 cells, 3 experiments, 1294 structures) (Rings: p=0.0002; Arcs: p=0.0007, Lines: p=0.139, Undefined clusters: p=0.00007)*.

### GSDMD expression level affects the density, but not the shape, of structures at the PM

Given the highly heterogeneous shape and stoichiometry of GSDMD pores in the plasma membrane, we wondered whether this is related to the availability of GSDMD during pyroptosis. Attenuated GSDMD transcription has been shown to prevent GSDMD-mediated pyroptosis [40, 41], but how reduced GSDMD expression affects GSDMD pore formation is unclear. To tackle this question, we explored pore formation at reduced GSDMD levels as compared to the relatively high levels obtained by the CMV promotor used in HEK293T DS cells. For this purpose, we stably expressed mGSDMD-mEGFP under regulation of a PGK promotor (HEK293T DS PGK cells) [42]. Analysis of mEGFP intensity of HEK293T DS PGK cells confirmed the reduced GSDMD expression compared with HEK293T DS cells (**Extended Figure 4A**). As expected, HEK293T DS PGK cells showed attenuated PM permeabilization and LDH-assayed cell lysis after pyroptosis induction (**Figure 5A-C**). We then performed DNA-PAINT analysis of the PSPMs generated from these cells (**Figure 5D and Extended Figure 4B**). Our analysis revealed a lower density of these structures at the PM (**Figure 5E and Extended Figure 4C**). Unexpectedly, however, neither the relative percentage of ring and arc structures nor their size was affected by the reduced expression levels (**Figure 5F-H**). This observation suggests that the heterogeneity of pore shapes and stoichiometries results from the assembly process rather than the availability of GSDMD, implying that functional determinants in the PM may be responsible. Taken together, these data suggest that the number of GSDMD pores on the PM is a critical factor in the cell death decision. This provides a rationale for the observations that pyroptosis can be rescued by eliminating GSDMD pores through appropriate membrane repair mechanisms [23, 24].

**Figure 5.**
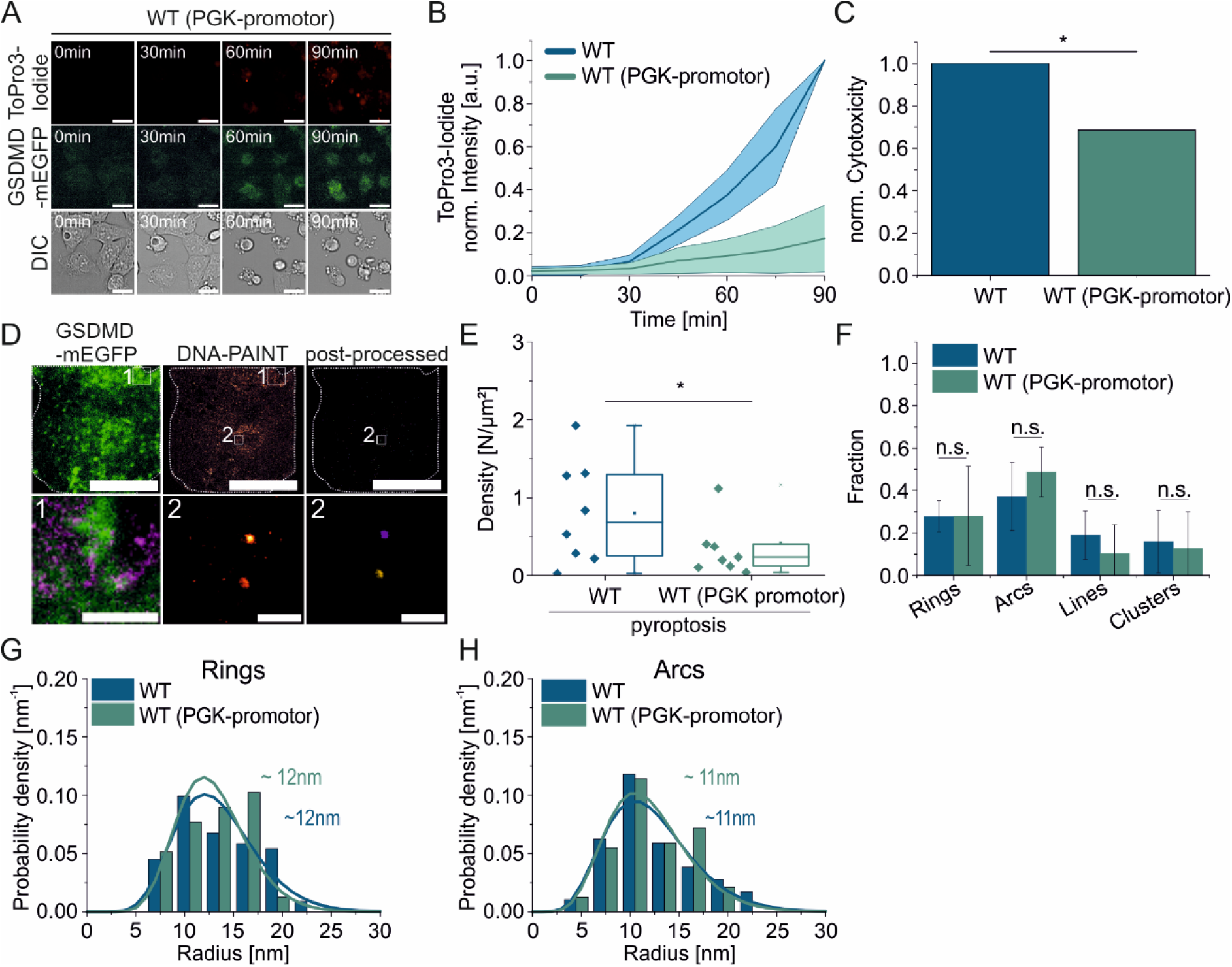
GSDMD expression level regulates the density of GSDMD structures in the PM. *(A) Representative confocal images of pyroptosis induction in HEK293T cells expressing mGSDMD-mEGFP with a PGK-promotor resulting in a lower expression level (green). Pyroptosis is monitored by ToPro3-Iodide staining (red) and morphological changes (DIC). Scale bars 20µm*. *(B) Quantification of PM permeabilization for HEK293T cells expressing mGSDMD-mEGFP with or without a PGK-promotor by normalized fluorescence intensity of ToPro3-Iodide (n= 3 experiments with >20 cells analyzed per experiment). Lines in the graph correspond to the average values from all measured cells and colored areas to data variability (mean ± SD*). *(C) Normalized cytotoxicity of HEK293T cells expressing mGSDMD-mEGFP with and without a PGK-promotor after 90 min of pyroptosis induction monitored by LDH-release assay (n=5 experiments, p=0.036)*. *(D) Upper row: overview of mGSDMD-mEGFP (PGK) (left) and DNA-PAINT localizations before (middle) and after (right) post-processing (Scale bars 20µm). Lower row: Correlation of mGSDMD-mEGFP (PGK, green) and DNA-PAINT localizations (magenta) from zoom-in area 1 in the upper row (left, scale bar 5 µm), zoom in of DNA-PAINT localizations from area 2 in the upper row before (middle) and after (right) post-processing. Scale bar 200 nm*. *(E) Density of super-resolved GSDMD structures after post-processing of DNA-PAINT localizations in HEK293T cells expressing mGSDMD-mEGFP with or without PGK-promotor (GSDMD WT n=8 cells; 3 experiments, GSDMD WT (PGK) n=9 cells; 3 experiments, p=0.169)*. *(F) Comparison between the relative distribution of the different GSDMD structure types in HEK293T cells expressing mGSDMD-mEGFP WT (n=8 cells, 3 experiments, 493 structures) or GSDMD-mEGFP WT (PGK) (n=9 cells, 3 experiments, 480 structures) (Rings: p=0.954; Arcs: p=0.231, Lines: p=0.346, Undefined clusters: p=0.927)*. *(G) Quantification of the relative distribution of radius of mGSDMD-mEGFP WT ring-like structures with and without PGK-promotor in PSPMs of pyroptotic HEK293T cells. Rings of mGSDMD-mEGFP WT (PGK) have an average radius of 12nm (Rings (PGK) n=30, n=9 cells; 3 experiments)*. *(H) Quantification of the relative distribution of the radius of mGSDMD-mEGFP WT arc-like structures with and without PGK-promotor in PSPMs of pyroptotic HEK293T cells. Arcs of mGSDMD-mEGFP WT (PGK) have an average radius of 11nm (Arcs (PGK) n=67, n=9 cells; 3 experiments)*.

### Phosphoinositide metabolism regulates pyroptosis

Harvesting the potential of PSPMs to monitor GSDMD nanostructures directly at the PM of pyroptotic cells, we reasoned that the PM nano-environment may regulate GSDMD pores. Several studies showed that phosphoinositides, especially PI(4,5)P_2_, play an important role in the permeabilization activity of GSDMD [7, 22, 28, 29]. Therefore, we monitored PI(4,5)P_2_ metabolism during pyroptosis and evaluated its effects on GSDMD structures and density at the PM. To this end, we co-expressed the PI(4,5)P_2_ sensor PH-PLCδ1-iRFP [43, 44] and the PI(3,4,5)P_3_ sensor PH-Akt-EGFP [45, 46] together with GSDMD-mCherry in HEK293T Cas1 cells and monitored the permeabilization of pyroptotic cells by SytoxBlue staining (**Figure 6A**). In the early stage of pyroptosis (approximately 10-20 min), we observed a gradual shift of the PH-PLCδ1-iRFP sensor from the PM to the cytosol, indicating a decrease of PI(4,5)P_2_ at the PM during pyroptosis (**Figure 6A-C**). Intriguingly, the decrease in PI(4,5)P_2_ signal at the PM correlated with a slight increase in PI(3,4,5)P_3_ levels at the early time points after pyroptosis induction (**Figure 6A-C**). This suggests that conversion of PI(4,5)P_2_ to PI(3,4,5)P_3_ occurs at an early stage of pyroptosis, which precedes detectable permeabilization of the PM, as shown by SytoxBlue staining (**Figure 6A**). To further investigate the importance of PI(3,4,5)P_3_ at the PM during pyroptosis, we inhibited PI(3,4,5)P_3_ production in pyroptotic cells by treatment with the PI3-kinase inhibitor Wortmannin (Wtm) [47]. Treatment with Wtm resulted in a complete loss of PI(3,4,5)P_3_ levels in the PM (**Extended Figure 5A**). Strikingly, cells lacking PI(3,4,5)P_3_ showed a significant decrease and delay in cell permeabilization after pyroptosis induction (**Figure 6D and Extended Figure 5C, D**). Treatment with Wtm without pyroptosis induction was well tolerated by the cells and we did not observe any visible changes in cell morphology or membrane permeabilization (**Extended Figure 5C and E**). However, because of the potential cell toxicity, we alternatively inhibited PI(3,4,5)P_3_ production by overexpressing a membrane-bound version of the PI(3,4,5)P_3_ phosphatase Pten [48] (**Extended Figure 5B**). Similar to Wtm treatment, overexpression of Pten resulted in a significant decrease in cell permeabilization upon pyroptosis induction (**Figure 6E and Extended Figure 5F-H**). Overall, these results demonstrate a critical role of PI(3,4,5)P_3_ in regulating membrane permeabilization during pyroptosis.

**Figure 6.**
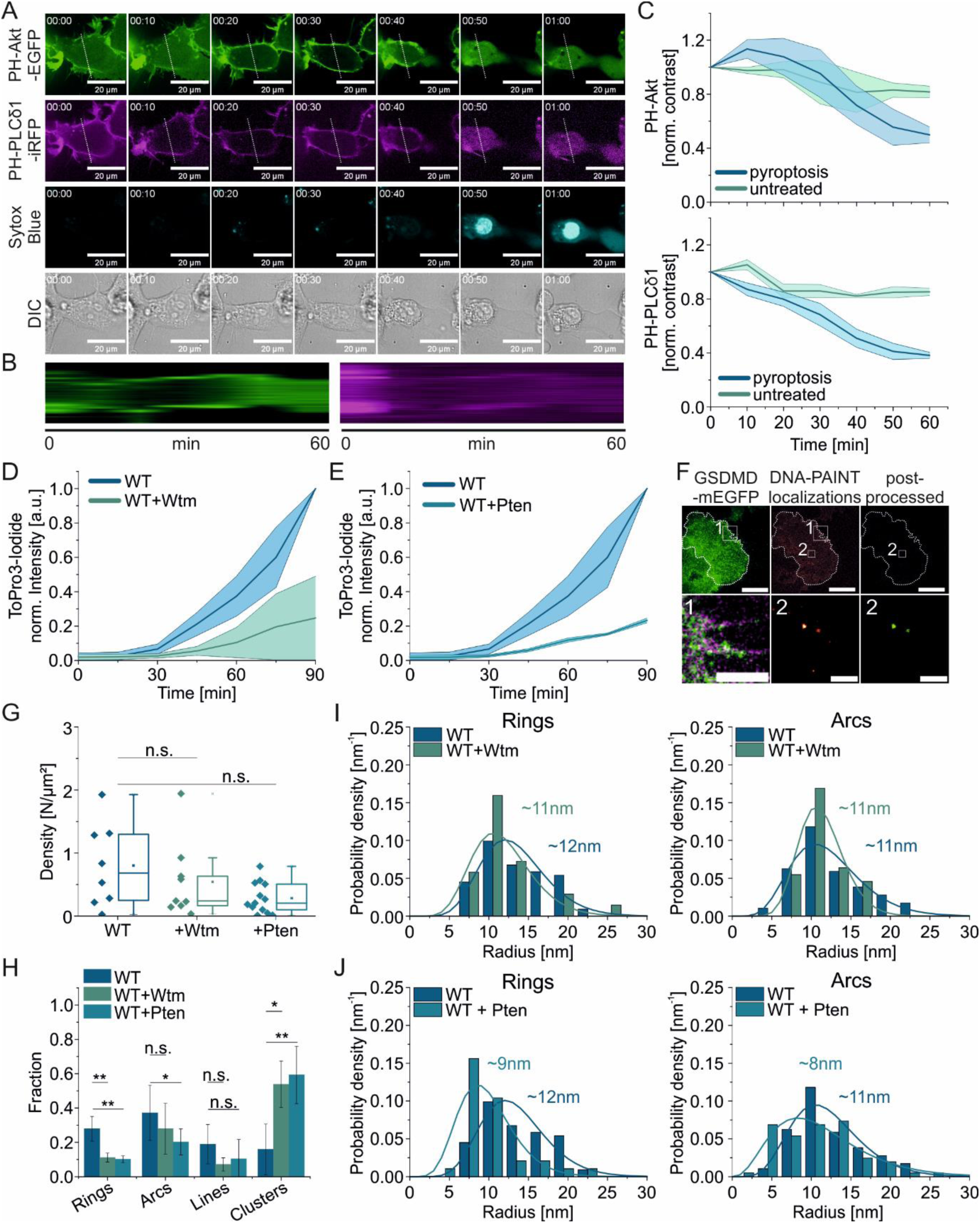
PI(3,4,5)P_3_ facilitates PM permeabilization in pyroptosis and the formation of GSDMD functional structures with an increased radius. *(A) Representative confocal images of pyroptosis induction for 60 min and PI(4,5)P_2_ and PI(3,4,5)P_3_ conversion in HEK293T cells expressing DmrB-mCas11, GSDMD-mCherry, the PI(4,5)P_2_ marker PH-PLCδ1-iRFP (magenta) and the PI(3,4,5)P_3_ marker PH-Akt-EGFP (green). Pyroptosis induction is monitored every 10 min by nuclear staining with SytoxBlue (blue) and morphological changes (DIC)*. *(B) Representative kymographs of PH-Akt-EGFP (green) and PH-PLCδ1-iRFP (magenta) over 60min of pyroptosis induction along the line indicated in (A)*. *(C) Normalized contrast (PM/Cytosol) of PH-Akt-EGFP (or PH-Akt-mScarlet) (upper panel) and of PH-PLCδ1-iRFP (lower panel) in untreated or pyroptotic HEK293T cells over 60 min monitoring (3 experiments with > 5cells)*. *(D) Quantification of PM permeabilization of HEK293T cells expressing mGSDMD-mEGFP with or without treatment with the PI3-Kinase inhibitor Wtm (10µM) by normalized fluorescence intensity of ToPro3-Iodide (n= 3 experiments with >20 cells analyzed per experiment). Lines in the graphs in C and D correspond to the average values from all measured cells and colored areas to data variability (mean ± SD)*. *(E) Quantification of PM permeabilization of HEK293T cells expressing mGSDMD-mEGFP with or without expression of the PI(3,4,5)P_3_-phosphatase Pten by normalized fluorescence intensity of ToPro3-Iodide (n= 3 experiments with >20 cells analyzed per experiment)*. *(F) Upper row: overview of mGSDMD-mEGFP and DNA-PAINT localizations before and after post-processing of a PSPM from a pyroptotic cell with Pten overexpression (Scale bars 20µm). Lower row: Correlation of mGSDMD-mEGFP (green) and DNA-PAINT localizations (magenta) from zoom-in area 1 in the upper row (left, scale bar 5 µm), zoom in of DNA-PAINT localizations from area 2 in the upper row before (middle) and after (right) post-processing. Scale bar 200 nm*. *(G) Density of super-resolved GSDMD structures after post-processing of DNA-PAINT localizations in HEK293T cells expressing mGSDMD-mEGFP with or without treatment with 10µM Wtm or expression of Pten (GSDMD WT n=8 cells; 3 experiments, +Wtm n=7 cells; 3 experiments, p=0.415; +Pten n=12 cells, 4 experiments, p=0.065)*. *(H) Comparison between the relative distribution of the different GSDMD structure types in HEK293T cells expressing mGSDMD-mEGFP WT (n=8 cells, 3 experiments, 493 structures) with treatment of 10µM Wtm (n=7 cells, 3 experiments, 385 structures) or with expression of Pten (n=8 cells, 3 experiments, 594 structures); (+Wtm: Rings: p=0.003; Arcs: p=0.389; Lines: p=0.174; Undefined clusters: p=0.012); (+Pten: Rings: p=0.002; Arcs: p=0.018; Lines: p=0.324; Undefined clusters: p=0.002)*. *(I) Quantification of the relative distribution of the radius of mGSDMD-mEGFP WT ring-and arc-like structures with or without 10µM Wtm in PSPMs of pyroptotic HEK293T cells. Wtm treted cells have rings with an average radius of 11nm (n=23) and arcs of 11nm (n=73)*. *(J) Quantification of the relative distribution of the radius of mGSDMD-mEGFP WT ring-and arc-like structures with or without Pten expression in PSPMs of pyroptotic HEK293T cells. Pten expressing cells have rings with an average radius of 9nm (n=32) and arcs of 8nm (n=68)*.

### PI(3,4,5)P_3_ is critical for the formation of functional GSDMD structures in the PM

To further elucidate the role of PI(3,4,5)P_3_ in membrane permeabilization by GSDMD pores, we performed DNA-PAINT on PSPMs from pyroptotic cells with reduced PI(3,4,5)P_3_ levels. Depletion of PI(3,4,5)P_3_ by either overexpression of Pten or treatment with Wtm, did neither affect the formation of GSDMD puncta after pyroptosis induction (**Figure 6F and Extended Figure 5I-K**) nor the density of GSDMD structures at the PM (**Figure 6G**). However, it led to a significant impairment of the formation of ring-shaped GSDMD assemblies and a strong attenuation of the number of arc-like GSDMD structures, compared to pyroptotic HEK293T DS WT cells (**Figure 6H**). Notably, ring structures in Wtm-mediated PI(3,4,5)P_3_-depleted cells were smaller than rings formed in the presence of PI(3,4,5)P_3_ (∼11 nm compared to ∼12 nm; **Figure 6I**). This effect was even more pronounced in Pten-overexpressing cells, where both the radius of arcs and rings were smaller (with an average radius of ∼8 and ∼9 nm, respectively; **Figure 6J**). Under both conditions, the size of the rings were affected, indicating a decisive role of PI(3,4,5)P_3_ in yielding the final, fully assembled GSDMD ring structures (**Figure 6I and J**).

### PI(3,4,5)P_3_ stabilizes GSDMD pores in the PM

To rationalize the severely hampered GSDMD pore formation in PI(3,4,5)P_3_-depleted cells, we developed a molecular dynamics (MD) simulation system. In a previous study, we identified highly positively charged clusters of arginine and lysine residues on both sides of human GSDMD that interact with PI(4,5)P_2_ and hypothesized that the high negative charge of PIPs might stabilize this interface and hence facilitate GSDMD assembly [28]. To recapitulate the behavior of PI(3,4,5)P_3_ in the pore formation of GSDMD, we simulated 16-meric half-rings of the structurally available human GSDMD embedded in a DOPC membrane with either PI(4,5)P_2_, PI(3,4,5)P_3_, or no PIPs in the interfaces between adjacent subunits. The choice of a simulation environment with a specific lipid composition (where no other lipids can interfere with PIPs) and with PIPs at GSDMD subunit interfaces was dictated by the need to have fully defined conditions to specifically answer the question of how PI(3,4,5)P_3_ compares to PI(4,5)P_2_ in stabilizing GSDMD pores. In this setup, an open membrane edge formed after the lipid bilayer detached from the GSDMD arc [28]. The associated line tension placed the arc under a force of about 44 pN [49]. Because of this large mechanical tension, the interfaces in the rings cracked within the first few hundred nanoseconds of our MD simulations (**Extended Videos 1-3**). Without PIPs in the interfaces, the half-rings typically cracked at 4-5 sites and the assembly equilibrated as a round pore (**Figure 7A, C**). The presence of PIPs in the interfaces reduced the number of cracks to 1-3 with PI(4,5)P_2_ and even further to 1-2 when PI(3,4,5)P_3_ was present, indicating a lipid charge-dependent stabilizing effect. In addition, the shape of the pores with PIPs was often more asymmetric with a membrane edge still evident. The increased stability of the assemblies containing bridging PIPs was evident from the approximated free energy of cracking (**Figure 7B**). Compared with the assemblies without PIPs, in the systems with PI(4,5)P_2_, the distances between the centers of mass of adjacent subunits were more stable even below 28 Å, which broadened the free energy minimum. With bridging PI(3,4,5)P_3_, the distance between two adjacent subunits was more stable maintained around 27.5 Å, resulting in a narrower free energy minimum and a greatly increased barrier to cracking. As such, with PI(3,4,5)P_3_ bound to the interface, ‘near cracking’ distances (∼30 Å) between two neighbors were reached less frequently.

**Figure 7.**
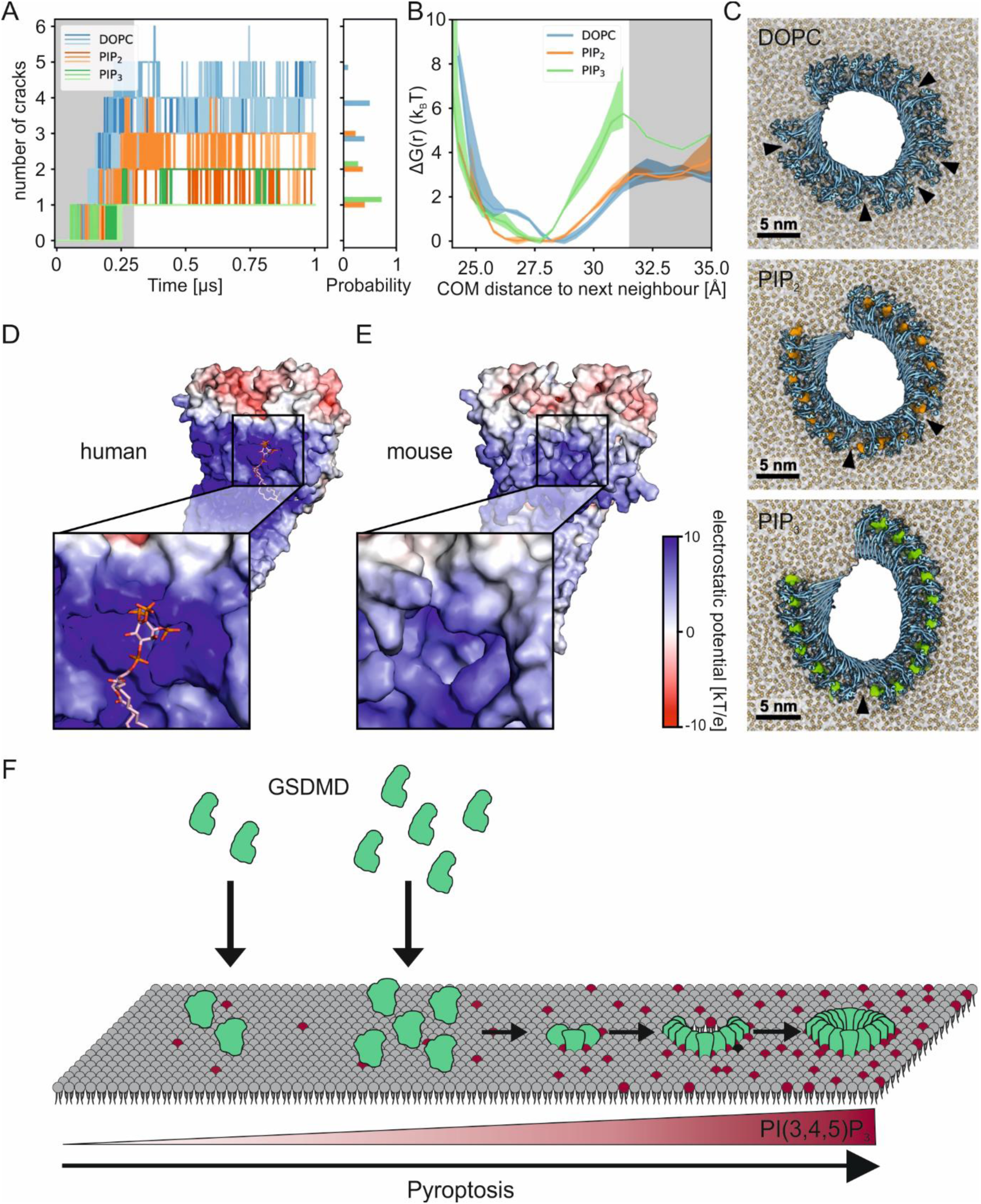
PI(3,4,5)P_3_ stabilizes the interface between GSDMD subunits. *(A) Number of cracks over the course of the 1 µs long simulations shown for each of the three replicates of the system without PIPs (shades of blue), the system with PI(4,5)P_2_ (shades of orange), and the system with PI(3,4,5)P_3_ (shades of green) shown on the left and the probability distribution of the number of cracks during the last 700 ns shown on the right (one color per system). The shaded area shows the equilibration phase that was excluded from the density*. *(B) Free energy of cracking estimated from free energy profile as function of the center-of-mass (COM) distance between neighboring subunits. The average of the three replicates is shown as a solid line (colored shading: SEM). Rare cracking events in the PI(3,4,5)P_3_ system prevent reliable error estimates in the cracked region* (*r* > 31.5 *Å; shaded area)*. *(C) Representative snapshots of the GSDMD assemblies after 1 µs of unbiased simulation without PIPs (top), with PI(4,5)P_2_ (middle), and with PI(3,4,5)P_3_ (bottom) in top view. The protein is shown in blue cartoon representation. PI(4,5)P_2_ and PI(3,4,5)P_3_ molecules are shown as orange and green densities, respectively. Black triangles show cracks in the backbone of globular domains of the assembly*. *(D) Electrostatic potential surface of a hGSDMD dimer calculated with APBS^1^ shown with a PI(3,4,5)P_3_ molecule bound to the interfacial binding site. This PI(3,4,5)P_3_ was not included in the electrostatic potential calculation*. *(E) Electrostatic potential surface of a mGSDMD dimer modelled with AlphaFold*. *(F) Model of GSDMD pore regulation in the PM: GSDMD concentration affects the density but not the assembly process, suggesting that pore formation is probably an autocatalytic process, i.e. the growing of the pore is faster than the formation of new seeding points for pore formation. The assembly process is modulated by the membrane environment. Pore widening would decrease due to line tension, but PI(3,4,5)P_3_, which levels increase during pyroptosis, provides a transient stabilization that prolongs pore growth*.

We further characterized the electrostatic environment around GSDMD. At the interface, the charge of the basic clusters mentioned above is not sufficiently balanced by acidic protein residues. Therefore, like the entire bottom of GSDMD, the interface region between two subunits exhibits a strong positive electrostatic potential (**Figure 7D**). The alphafold-modelled [50, 51]mouse structure presents a similar environment (**Figure 7E**). We also analyzed the specific binding modes exhibited by different PIPs in the non-cracked binding site (**Extended Figure 6**). In a previous study, we identified lateral binding sites of human GSDMD comprising five charged residues (K43, R53, K55, R153, and K235) interacting with PI(4,5)P_2_. Driven by electrostatics, both the PI(4,5)P_2_ and PI(3,4,5)P_3_ lipids, with their four and six negative charges, respectively, interact with the highly positively charged interface. However, unlike PI(4,5)P_2_, PI(3,4,5)P_3_ can accommodate the charges of all five residues simultaneously. These extensive interactions lead to an increase in contacts between PI(3,4,5)P_3_ and K43, R153, and K235 compared to PI(4,5)P_2_ (**Extended Figure 6**). We propose that this increase in charge density of PI(3,4,5)P_3_ compared to PI(4,5)P_2_ is capable to electrostatically stabilize the interface sufficiently to compensate the cracking force caused by the line tension. In turn, this may stabilize the opening of GSDMD pores, thus favoring the growth of arc and ring GSDMD structures (**Figure 5**).

## Discussion

Here, we report the first structural insight into GSDM pores at the PM of pyroptotic cells and reveal a novel function of the lipid environment as a master regulator of pyroptosis through PIPs-mediated stabilization of functional GSDMD structures. Based on our observations, we propose a model for PM-mediated regulation of pyroptosis through modulation of both, GSDMD pore concentration and functionality. This mechanism is tightly regulated by the PIP metabolism at the PM, with PI(3,4,5)P_3_ promoting the pore assembly process (**Figure 7F**). Importantly, our work establishes a correlation between pyroptosis outcome and the number and architecture of GSDMD structures as we directly evaluate them in their native PM environment by using our newly established PSPM technology.

Until now, the resolution of GSDMD pores in cells has been hampered by the drastic morphological changes of cells undergoing pyroptosis causing cell swelling and partial detachment from the cell support, as well as by the strong fluorescent background of cytosolic labeled-GSDMD. To overcome these problems, we have developed PSPMs that allow the fabrication of an immobilized, flat and integral PM sheets for clear visualization of GSDMD structures at the single pore level. Compared to existing methods for “cell-unroofing”, PSPMs offer the additional advantage of tethering the PM to a polymer support. This approach allows adherence of pyroptotic cells to the substrate and thus imaging of GSDMD structures at the PM unbiased by local membrane curvatures and different distances from the polymer support. In addition, PSPM production does not require harsh and lengthy treatments, e.g., with hypotonic solutions or freeze-fracture techniques [52-55] and can be easily performed in less than 5 min directly at the microscope. Like polymer-supported artificial membranes [56, 57], the cushion formed by the polymer between the PM and the solid support avoids friction of the membrane with the glass surface. Similar to other methods for isolating PM sheets, this method affects the curvature of the membrane, and we cannot rule out the possibility that this may affect protein localization and function. However, using this procedure, farnesylated proteins in PSPMs and in intact cells show similar diffusion behavior (**Extended Figure 1** and [36]). In general, PSPMs can be used to study protein diffusion and assembly in the native PM environment. Importantly, this approach makes the cytosolic side of the PM accessible for efficient labeling of all membrane-associated structures, including endogenous proteins, and for investigation by various microscopy techniques, such as AFM and SEM.

By combining PSPMs with super-resolution DNA-PAINT, we were able to resolve nanoscopic structures with a resolution of less than 10 nm. GSDMD pores at the PM exist as highly heterogeneous structures in shape, size, and stoichiometry. Our data in cells are consistent with previous structural studies of GSDMD assemblies in artificial membranes [7, 11, 22] and with the structural heterogeneity exhibited by other pore-forming proteins [58, 59]. We exclude that PSPM production had a significant impact on GSDMD structures because disruption of the cytoskeleton is a feature of pyroptotic cells [60] and Latrunculin B treatment occurred only after pyroptosis induction and GSDMD pore formation. Furthermore, changes in membrane curvature due to membrane tethering did not prevent the formation of functional GSDMD structures, as the timing of cell death was similar in tethered and untethered cells (**Figure 1F and G**).

Furthermore, we have presented here a compelling approach to accurately quantify the stoichiometry of protein complexes directly in their native PM environment. To quantify the stoichiometry of GSDMD pores in the PM, we performed qPAINT analysis by developing an orthogonal surface functionalization system for simultaneous immobilization of DNA origami and PSPMs having GSDMD pores. Our analysis revealed a broad distribution of stoichiometry of GSDMD assemblies in the PM with an average stoichiometry of 14 subunits for arcs and 18 for rings. These numbers agree well with the stoichiometry of arcs but not of rings previously measured by AFM in supported bilayers (16 and 30 subunits, respectively [22]). While we cannot rule out underestimation in subunit counts due to measurement conditions, such as inaccessibility of the anti-GFP nanobody due to steric hindrance at the pore or overestimation of the labelling efficiency of the nanobody [61], we found that our stoichiometry values for rings have a similar distribution to the slit shapes described in [22] (with an average of 21 subunits). In our analysis, however, we could not identify slit structures, which could be an explanation for the lower stoichiometry observed for rings. Another factor could be the different environment provided by the native PM compared to artificial systems that can actively control the equilibrium of pore formation, leading to the formation of smaller ring structures that shift the overall distribution to lower stoichiometry values.

Of all detected structures, ring-and arc-shaped architectures are probably most relevant to the execution of pyroptosis by PM permeabilization, because the dysfunctional mGSDMD-C192A mutant assembled predominantly in small oligomers (termed undefined clusters), but not in ring-and arc-shaped structures (**Figure 4**). Remarkably, this mutant affects higher-order oligomerization but does not abolish membrane binding. Because Cys192 still allows formation of small structures, we speculate that the reduction in formation of arcs and rings is due either to reduced protein recruitment to the PM mediated by Cys192 or to a Cys192-mediated interaction between small oligomers which serve as building blocks in the formation of GSDMD pores. Whether low-order oligomers may already be functional for the passage of ion molecules, as previously proposed [28, 39], requires further investigation.

Importantly, our methodological approach offers the unique advantage of investigating how regulatory mechanisms of the PM environment influence pyroptosis through modulation of GSDMD pores. Our analysis of the correlation between the outcome of pyroptosis and the density of GSDMD structures at the PM provides a rationale for the idea that membrane repair mechanisms rescue cells from death by removing GSDMD pores at the PM [23, 24]. Finally, we detected a change in PM lipid composition, namely a conversion of PI(4,5)P_2_ to PI(3,4,5)P_3_, at early stages of pyroptosis induction that precedes PM permeabilization. It is known that the PI3-kinase, which is responsible for PI(3,4,5)P_3_ synthesis, can be activated by Ca^2+^ influx [29, 62, 63]. It remains to be elucidated what triggers this process and whether it occurs upstream of GSDMD pore formation or whether it is mediated by early Ca^2+^ influx through small GSDMD oligomers.

Remarkably, PI(3,4,5)P_3_ plays a functional role in pyroptosis, as cells with reduced PI(3,4,5)P_3_ levels exhibited a significant reduction in PM permeabilization. Previous studies have shown that negatively charged lipids, particularly PI(4,5)P_2_, play a critical role in GSDMD pore formation [7, 22]. More recently, electrophysiological measurements have shown that GSDMD pores are highly dynamic multimolecular structures whose function is controlled by Ca^2+^-dependent local PI(3,4,5)P_3_ concentrations [29]. Here we demonstrate that PI(3,4,5)P_3_ is able to stabilize GSDMD assemblies more than PI(4,5)P_2_, both in experiments and in molecular dynamics simulations, and that an increase in PI(3,4,5)P_3_ concentration in the PM during pyroptosis is essential for the stabilization of the fully assembled functional GSDMD pores. Because the GSDMD density at the PM is not affected by the lack of PI(3,4,5)P_3_, we hypothesize that PI(4,5)P_2_ is sufficient to target GSDMD to the PM but results in smaller and mechanically weaker assemblies.

In summary, our work provides mechanistic insight into the formation and regulation of GSDMD pores on the PM during pyroptosis. We show that oligomerization of GSDMD leads to various distinct structures with different permeabilization capabilities. The formation and stabilization of functional pores is regulated by cells through changes in PM lipid composition. The molecular basis for this effect results from the ability of the highly negatively charged PI(3,4,5)P_3_ to form strong electrostatic interactions with GSDMD. Our work opens the possibility of modulating GSDMD pore opening by PIP metabolism to control pyroptosis in disease.

## Material & methods

### Cell culture

HEK293T stable DmrB-mCas1 (Tet-On vector) and double-stable DmrB-mCas1 (Tet-On) + mGSDMD-N-mEGFP-C (with mEGFP just before the caspase cleavage site, as in [33]) were cultured at 37°C and 5% CO_2_ MEM Eagle (PAN Biotech, P04-09500) supplemented with 10% Tet-On system approved FBS (Gibco, A4736401), 1% non-essential amino acids (MEM NEAA, PanBiotech, P08-32100) and 1% HEPES buffer 1M (PanBiotech, P05-01100). Cells were transfected at 80% confluence by Calcium-phosphate precipitation over-night. The day before microscopy, cells were detached by treatment with Trypsin/EDTA (Capricorn Scientific, TRY-1B10) at RT and seeded on the respective microscopy supports.

### Pyroptosis induction and permeabilization kinetics

16-24 hours before the experiments, cells were treated with 500ng/ml Doxycycline (Sigma Aldrich, D3447) to induce expression of DmrB-mCas1. Pyroptosis was induced by addition of 500nM B/B-Homodimerizer (Takara Bio, AP20187) at 37°C and 5% CO_2_. To observe pyroptotic cell permeabilization, the sample was treated with 0.5 µM To-Pro-3 Iodide (Thermo Fisher Scientific, T3605) or 3 µM SytoxBlue Cell Death stain (Thermos Fisher Scientific, S34857). Pyroptosis kinetics measurements were performed by acquiring images before pyroptosis induction and afterwards with intervals of 15 min at a Spinning disk confocal microscope (CellVoyager^TM^ CQ1 Benchtop High, Yokogawa) at 37 °C with a dry 40x objective.

### Polymer-supported PMs

In order to generate polymer-supported plasma membranes (PSPMs), gridded coverslips (Ibidi, 10817) were intensively cleaned with isopropanol before and after plasma cleaning (Plasma Cleaner femto 1A, Diener electronics) at 100% output power for 15 min. Subsequently, the slides were coated with a 30/70% (w/w) mixture of poly-L-lysine coupled to a polyethylene glycol functionalized with either a HaloTag-ligand (PLL-PEG-HTL) and an RGD-peptide (PLL-PEG-RGD), respectively [64]. Cells for PSPM synthesis were transfected with pDisplay-HaloTag-mTagBFP-TMD-GSlinker (TMD sequence: ASALAALAALAALAALAALAALAKSSRL) and seeded on the functionalized slides the day before microscopy. While the RGD-peptide functionalization allows attachment of the cells by integrin interactions, the covalent HTL-HaloTag interaction allows stable tethering of the PM to the surface. To minimize side effects due to membrane anchoring, the percentage of HaloTag-ligand binding PM on the polymer-coated surface was kept at 30%. For final PSPM generation, cells were treated with 10µM Latrunculin B (abcam, ab144291) for 5 min at 37°C and subsequently the cell body was removed by sheer forces through heavy pipetting directly at the microscope. Cell debris was removed by washing the sample 3x with 1ml PBS (DPBS, PanBiotech, P04-35500). Afterwards, PSPMs were fixed with 4% paraformaldehyde.

### Scanning electron microscopy

For scanning electron microscopy (SEM), PSPMs of HEK393T DS cells were prepared as described above. After removal of the cell body, PSPMs were fixed with 2.5% glutaraldehyde (Science Services, Germany) in 0.2 M HEPES buffer pH 7.2 (Roth, Germany) for 30 min at room temperature, then washed twice with 0.2 M HEPES buffer pH 7.2, and once with Milli-Q water (Merck Millipore). Samples were stained with aqueous 0.1% uranyl acetate (Science Services) for 20 minutes, washed once with Milli-Q water (Merck Millipore), dehydrated in a graded ethanol series of 50%, 70%, 80%, 90% and twice in 100% ethanol for four minutes each. Finally, PSPMS were critical point dried in 100% ethanol in a critical-point dryer (Leica CPD300, Leica, Austria) with liquid carbon dioxide as transition fluid, and then glued onto Leit-tabs (Plano, Germany) mounted on aluminum stubs (Plano, Germany) and sputter-coated with a 3 nm thin gold layer (ACD600, Leica, Austria). SEM images were acquired with a Zeiss Auriga FEG-SEM (Zeiss, Germany) operating at an accelerating voltage of 4kV, with an InLens detector at 4.2 mm working distance.

### Total internal reflection fluorescence (TIRF) microscopy

All microscopy imaging experiments were performed at an inverted Olympus IX-81 microscope equipped with a motorized quad-line TIR-illumination condenser (cellTIRF-4-Line, Olympus), a motorized xy-stage (IM 120×80, Märzhäuser), a 100x oil immersion objective (UAPON 100x TIRF, NA 1.49, Olympus), a large incubator with temperature control (TempController 2000-2, CellVivo) and a CO_2_-controller (CO_2_-controller 2000, CellVivo). In order to ensure PM tethering by HaloTag anchor expression, mTagBFP excitation was achieved by a 405 nm diode laser (Olympus), GSDMD-mEGFP oligomer formation was monitored by excitation with a 488 nm diode-pumped solid-state laser (Olympus) and Pten expression was ensured by excitation of mScarlet with a 561 nm diode-pumped solid-state laser (Olympus), both set to TIR conditions. Cell permeabilization was monitored by nuclear staining by the cell-impermeable dye To-Pro-3 Iodide via epi-mode excitation with a 640 nm diode laser (Olympus). Laser lines were filtered by clean-up filters (405nm: BrightLine HC 390/40, Semrock, 488nm: BrightLine HC 482/18, Semrock, 561nm: BrightLine HC 561/14, Semrock and 640nm: BrightLine HC 640/1, Semrock). Fluorescence emission was filtered by a bandpass filter (BrightLine HC 446/523/500/677) and, in addition, single bandpass filters (BrightLine HC 390/40, 482/18, 561/14 and 640/14, Semrock) for each channel, respectively before detection with a sCMOS camera (ORCAFlash 4.0 V3, Hamamatsu). Images were acquired with the software CellSens 3.2 (Olympus) with an exposure time of 32 ms and 2×2 pixel binning, resulting in a pixel size of 130 nm.

### FRAP & Single-molecule tracking

PSPM lipid mobility was investigated by FRAP and single-molecule tracking analysis at room-temperature. First, PSPM generation and TIRF microscopy were performed as described above using HEK293T cells stable expressing DmrB-mCas1 and mGSDMD-mEGFP and transiently expressing the HaloTag-mTagBFP-TMD anchor and Farnesyl-mCherry as a membrane marker. After PSPM generation directly at the microscope, a circular region with a radius of 5 µm was bleached using the 405 nm laser set to 100% for 10 sec. Immediately afterwards, fluorescence recovery of farnesyl-mCherry was detected over 200 sec with 1 sec intervals. The apparent diffusion coefficient was derived by the photo-bleaching corrected fluorescence recovery over time:

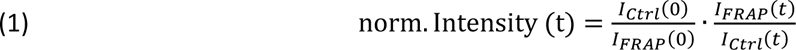

A mono-exponential fit provided the half-life τ_1/2_ which, together with the radius r, lead to the diffusion coefficient by the Soumpasis equation [65]:

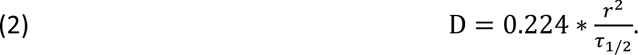

In order to analyze dynamics at single molecule level by tracking, we substoichiometrically labeled farnesyl-mCherry by 10 pM of a Dy647-conjugated LaM-anti-mCherry nanobody. After incubation for 10 min, diffusion of single farnesyl-mCherry molecules was monitored for 500 frames with a 30 Hz frame rate. The diffusion coefficient was obtained by MSD analysis using the software SLIMFast 4C [34] and trajectories were extracted using the software TrackIt [66].

### Atomic force microscopy

AFM experiments were performed on a JPK nanowizard atomic force system (JPK Instruments) mounted on an inverted Olympus IX-71 microscope (Olympus). PSPMs were imaged using silicon nitride cantilevers (SNL-10, Bruker) with a typical spring constant of 0.09 N/m in intermittent contact mode. Here, the cantilever oscillation was fine-tuned between 2-5 kHz and the amplitude set to 1 V. Imaging was performed at RT with a scan rate between 0.6 to 1 Hz. Height profiles where obtained using a smoothing function with the JPKSPM data processing (JPK instruments).

### DNA-PAINT

PSPMs were fixed after PSPM generation with 4 % PFA solution for 15 min at RT and subsequently washed 5x with 1 ml PBS (PanBiotech, P04-35500) before storage at 4°C. For DNA-PAINT preparation, the samples were reduced with 1 mg/ml sodium borohydride for 7 min at RT, washed 4x 5 min with 1ml washing buffer (Massive Photonics) before incubation with 50 µM of the anti-GFP nanobody + docking strand 3 (MASSIVE-TAG-Q anti-GFP DNA-PAINT Kit, Massive Photonics) dissolved in Antibody-incubation buffer (Massive Photonics) for 2 h at RT or at 4 °C over-night. Afterwards, the sample was washed 3x with 5ml washing buffer and incubated with a 1:1000 dilution of 90 nm gold nanorods (Cytodiagnostics, G-90-20) serving as fiducial markers for 5 min at RT and washed again with washing buffer. To ensure removal of unspecifically bound nanobodies, the sample was washed with imaging buffer (Massive Photonics) before supplementation with 1 nM Cy3b-conjugated imager strand-3 (MASSIVE-TAG-Q anti-GFP DNA-PAINT Kit, Massive Photonics) dissolved in imaging buffer. All DNA-PAINT experiments were performed at the setup described in the “TIRF microscopy” paragraph. The temperature was kept stable at 27 °C and buffer evaporation was prevented by humidification. The Imager was excited with a 561 nm laser adjusted to a power density of approx. 100 W/cm^2^ in the focal plane. For each recording, typically 40.000 frames were acquired with 200 ms exposure time and 2×2 pixel binning. During acquisition, the focus plane was stabilized with a hardware autofocus system (IX2-ZDC2, Olympus).

DNA-PAINT raw data sets were processed with the Picasso software suite [67]. For analysis, the last 30.000 frames of each dataset were considered to minimize drift and unspecific signals. First, DNA-PAINT signals were identified and localized in “Picasso Localize”. The Box size was set to 7 px and the Min. Net. Gradient was typically adjusted to 3.000-5.000 to filter unwanted background signals and signals with a weak signal-to-noise ratio. Photon conversion parameters were set as follows: EM gain: 1; Baseline: 400; Sensitivity 0.46; Quantum efficiency 0.72, pixel size: 130 nm. Single emitter signals were fitted with by integrated Gaussian fit. In “Picasso Filter”, the fitted signals were filtered for localization precision with a threshold of 0.05 px / 6 nm. In “Picasso render” processed datasets were drift-corrected, first, by cross correlation and further corrected with gold nanorod fiducials. Afterwards, localizations were linked with a max. number of dark fames according to the average mean length of the localizations and 0.05 px according to the localization precision threshold. Bright super-resolved structures were picked with a pick diameter of 0.5 px and filtered by trace for frequent sampling over all frames to select for specific GSDMD interactions. With the “pick similar” function, specific structures with a frequent sampling pattern were picked automatically with a STDEV of localization number of 1.0. For further removal of accumulations of unspecific localizations, we filtered all picked structures by the number of localizations. Picked structures with less than 30 localization/30 000 frames were removed as according to DNA-Origami experiments with the same Imaging conditions, monomeric binding sites should exhibit this localization frequency in this range. From the remaining structures 50 random overview images per dataset were saved with a Zoom of 100 (Pixel size of 1.3nm) as 8-bit tiffs for structural analysis with ASAP (see below).

### Structural classification of super-resolved structures

Overview images of DNA-PAINT super-resolved GSDMD structures were analyzed for structural classification with the software Automated Structures Analysis Program (ASAP, [37]). First, structures were identified by “Connectivity” with the following parameters: Thresholding method: fixed; threshold multiplier: 0.1; no cleaner, Identification range size: 200-10000. The exported identification files with a pixel size of 1.3 nm (Zoom 100, see “DNA-PAINT”) were analyzed by radial profiling with a max. ring size of 50 px and no operations were selected. Based on observations on the appearance of GSDMD structures, the software was trained to classify all identified structures into the following shape classifications: Rings, Arcs, Lines and Undefined clusters. Training for automated structural classification was performed with the learner “discriminant” and by the descriptor “Raw Radial Profile”. Afterwards, all identified super-resolved GSDMD structures were automatically classified into the shapes according to the training process. Afterwards, classification was corrected if necessary in the module “classify”.

### qPAINT

Stoichiometric analysis by qPAINT requires a calibration with a known number of imager-binding sites to calculate the number of subunits in a GSDMD pore by the relative frequency of DNA-PAINT localizations [31]. We used single binding sites of DNA-Origami structures with 3×4 docking sites and a 20 nm mutual distance as a calibration for qPAINT analysis. DNA-Origami structures were assembled according to the protocol given in [67] using DNA-oligonucleotides designed with “Picasso design” [67]. In order to optimize the stoichiometric analysis, DNA-Origami structures were immobilized next to tethered PSPMs on an orthogonal functionalized surface. Orthogonal functionalization was achieved by micro-contact printing of PLL-PEG-HTL on plasma cleaned cover slides with PDMS stamps [64] creating square pattern of 100×100 µm in size, allowing tethering of PSPMs, and subsequent backfilling with PLL-PEG-Biotin for DNA-Origami immobilization. Cell seeding and incubation, pyroptosis induction and PSPM generation were performed as described above. For simultaneous DNA-PAINT imaging the sample was first prepared as described above and DNA-Origami immobilization was performed as described in [67], after incubation with the anti-GFP nanobody + docking strand 3 the sample was washed 2x with 500 µl buffer A+ following by incubation with 1 mg/ml Streptavidin (Streptavidin UltraPure, PanREAC AppliChem, A1495005) dissolved in buffer A+ for 5min. Unbound Streptavidin was washed out 2x with 500 µl buffer A+ and the buffer was exchanges by washing 2x with 500 µl buffer B+. 50 µl of a 1:10 dilution of purified DNA-Origamis in buffer B+ were supplemented for 30 min at RT and unbound molecules removed by washing with 500 µl B+. Afterwards, the Imager strand+Cy3b was used at 1 nM in buffer B+ for DNA-PAINT imaging. Imaging and DNA-PAINT post-processing was performed as described above but here the DNA-Origamis themselves served as fiducial markers. For qPAINT calibration, the average dark mean time of a single binding site for an individual experiment was gained by taking the localization frequencies of > 10 000 DNA-Origami single binding sites per experiment and fitting all dark mean times. For each structure classified as ring or arc the individual dark mean times were used to calculate the respective number of binding sites in this structure with the average dark mean time for a single binding site for each experiment. To obtain the number of GSDMD subunits in each structure the number of binding sites was corrected by the nanobodies labeling efficiency (Thevathasan, Kahnwald et al. 2019).

### Spinning Disk Confocal microscopy

For monitoring PI(4,5)P_2_ and PI(3,4,5)P_3_ dynamics during pyroptosis, Hek 296T cells stable expressing DmrB-mCas1 were transfected with pSems-GSDMD-mCherry, pcDNA3-AKT-PH-GFP or pSems-Akt-PH-mScarlet and pEGFP-iRFP-PH-PLCδ1 via Calcium-phosphate transfection over-night. A day before the experiment, the cells were seeded into 8-well plates (80821, ibidi) and treated with 500 ng/ml Doxycycline and directly before the experiment 3 µM SytoxBlue cell death stain (Thermo Fisher Scientific, S34857) is added. Pyroptosis was induced as described above. The cells were imaged before pyroptosis induction at every 10 min afterwards at a fully motorized inverted spinning disc microscope based on a Zeiss Cell Observer.Z1 and a CSU-X1 spinning disc unit (Yokogawa). The microscope was equipped with live cell imaging periphery based on a home-built incubation chamber enclosing the full microscope. Heating at 37°C is managed by a PID-controlled heater (The cube, Live Imaging Services). Samples were imaged at 37°C in a humidified 5% CO_2_ atmosphere (Zeiss CO2 module S1, Zeiss humidifier module S1). To monitor cell permeabilization, SytoxBlue was excited with a 405 nm diode laser (max. 50mW) and GSDMD-mCherry expression checked with a 561 nm diode laser (max. 40mW). While the lipid markers PH-PLCδ1-iRFP and PH-AKt-mEGFP were excited using a 488 nm optically pumped semiconductor laser (max. 100mW) and a 635 nm diode laser (max. 30mW), respectively, to monitor lipid dynamics. Emission light passed through a 63x oil immersion objective (Alpha Plan-Apochromat, NA 1.46) and a polychroic mirror and was filtered by a set of bandpass filters (BFP: 450/25, GFP: 525/25 mCherry: dualbandpass 500-554nm and 615-675nm and iRFP: 690/50) before being detected by a Hamamatsu ORCA Flash V3 resulting in a px size of 86 nm. The sample was kept stable during the experiment using a motorized xyz-stage (PZ-2000 XYZ, Applied Scientific Instrumentation) and a Definite Focus System (Zeiss). Every step was performed with the software Zeiss Zen 2.6. In order to analyze the contrast between the PM and the cytosol of the lipid markers during pyroptosis, the plasma membrane intensity and the intensity in the cytosol was measured using FIJI, the intensity ratio [PM/Cytosol] was calculated for both markers and each value normalized to the time point t=0.

### LDH assay

LDH-release cytotoxicity assay was performed in 96-well plates (Thermo Fisher Scientific, M33089) using the CyQUANT™ LDH Cytotoxicity assay (Invitrogen, C20301) according to the manufactures protocol. Each sample and control was performed in triplicates with 20 000 HEK293T cells stable expressing DmrB-mCas1 and mGSDMD-mEGFP (with or without PGK-promotor) seeded per well on the day before. The cells were treated with 500 ng/ml Doxycycline and pyroptosis induction was performed as described above. Fluorescence detection of LDH release was performed using the Tecon Plate reader Infinite 200 Pro M-Plex.

### Molecular dynamics simulations

We constructed a computational assay to quantify the stabilizing effect of PI(4,5)P_2_ and PI(3,4,5)P_3_ on the stability of GSDMD assemblies by adapting the MD simulation protocol of Schaefer & Hummer [28]. Specifically, we probed the resistance of GSDMD arcs with or without PIPs to mechanical force created by the line tension of the open membrane edge between the two ends of the arc.

Starting from the experimental structure of a complete hGSDMD ring (PDB Id: 6VFE [21]), we set up structures of 16-meric half-rings of hGSDMD in pore conformation and placed them in a 36 x 36 nm^2^ large membrane comprising 3750 DOPC lipids. We then placed a single PI(4,5)P_2_ or PI(3,4,5)P_3_ lipid with palmitoyl (16:0) and oleoyl (18:1) tails in each of the interfaces between neighboring subunits. We obtained PIP lipid topologies from CHARMM-GUI [68] and placed their headgroups into a previously described lateral binding site that involves K43, R53, K55, R135, and K235 [28]. For reference, we additionally performed three new replicate simulations of the previously described 16- meric hGSDMD half-ring in a pure DOPC membrane without any PIPs as set up and equilibrated previously [28]. We built the simulation systems as described in [28] and followed the same steepest descent energy minimization and equilibration protocols comprising three equilibration steps (5, 50 and 80 ns) with decreasing restraints on the atomic positions of the system. During the third equilibration simulation, the lipids (including the PIPs) were not subject to any restraints. This setup without other acidic lipid species and without a reservoir of PIP lipids in the bulk membrane was chosen specifically to quantify the stabilizing effect of different PIP species on the assembly in a controlled manner. For the bulk of the membrane, we used DOPC lipids because we found earlier that their high fluidity ensures relatively rapid dynamics of the membrane-inserted gasdermins [28].

After equilibration, we performed three production simulations with starting velocities drawn independently from the Maxwell-Boltzmann distribution for each of the systems. During these production simulations, no positional restraints were applied.

All production simulations were performed using Gromacs version 2022.4 [69]. The interactions were described with the CHARMM36m [70] force field and TIP3P water [71]. The MD simulations were performed at a constant pressure of 1 bar, maintained by using the semi-isotropic (X and Y dimensions coupled together) Parrinello-Rahman barostat [72] with a coupling time constant of 5 ps and a compressibility factor of 4.5 x 10^-5^ bar^-1^. To maintain a constant temperature of 37 ℃ we applied velocity rescale thermostats [73] separately to the protein, the lipids, and the solvent atoms every 1 ps. We computed electrostatic interactions using the particle-mesh Ewald (PME) algorithm [74] with a cut-off distance of 1.2 nm for the real space electrostatics. For van-der-Waals interactions, we used the same cut-off distance. Bonds to hydrogen atoms were constrained using the LINCS algorithm [75].

To analyze the effect of the presence or absence of PIPs on the stability of the GSDMD assembly, we measured the distance between the globular domains (residues 34-71, 123-159 & 215-239) of neighboring subunits. From the distribution p(r) of distances r between neighboring subunits we estimated the Gibbs free energy of GSDMD-interface cracking as

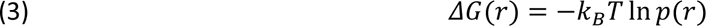

with *k_B_* Boltzmann’s constant and T the absolute temperature. For this reconstruction of ΔG, we only considered the last 700 ns of the replicate simulations. The constructed free energies were shifted vertically so that their global minimum, which corresponds to an intact GSDMD interface, is at zero. Based on the position of the maximum of the free energy landscapes, we considered a structure as cracked in the respective position if the distance r between two neighboring subunits exceeded 31.5 Å.

In addition, for neighboring subunits that do not have a cracked interface at a given time point, we also analyzed the PIP binding modes during the last 700 ns of the simulations. To do so, we counted the number of contacts of the respective bound PIP with all heavy protein atoms using a 3.6 Å cutoff and averaged them over all intact interfaces and over all considered frames. To calculate the electrostatic potential of GSDMD we used the APBS webserver [76]. For this, we uploaded a GSDMD dimer extracted from the last frame of one of the simulations with PI(3,4,5)P_3_ lipids and used the PDB2PQR tool to prepare input files for APBS. Since the input structure was the result of unbiased MD, we did not remodel any hydrogen atoms or optimized their bonding network. The APBS calculation itself was performed with default parameters.

Visual analysis and rendering were done using VMD [77], UCSF ChimeraX [78] and PyMOL [79].

### Material List

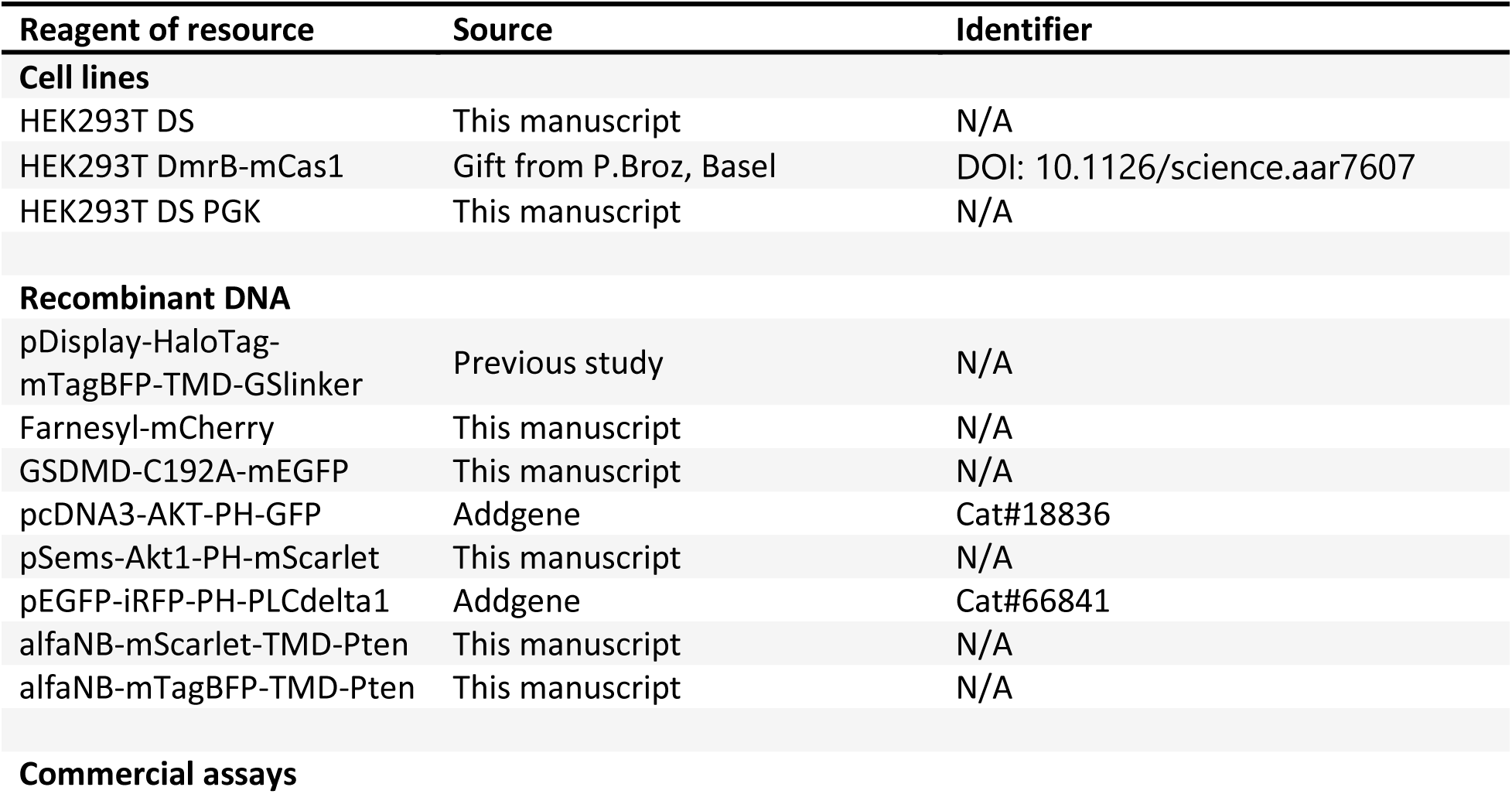
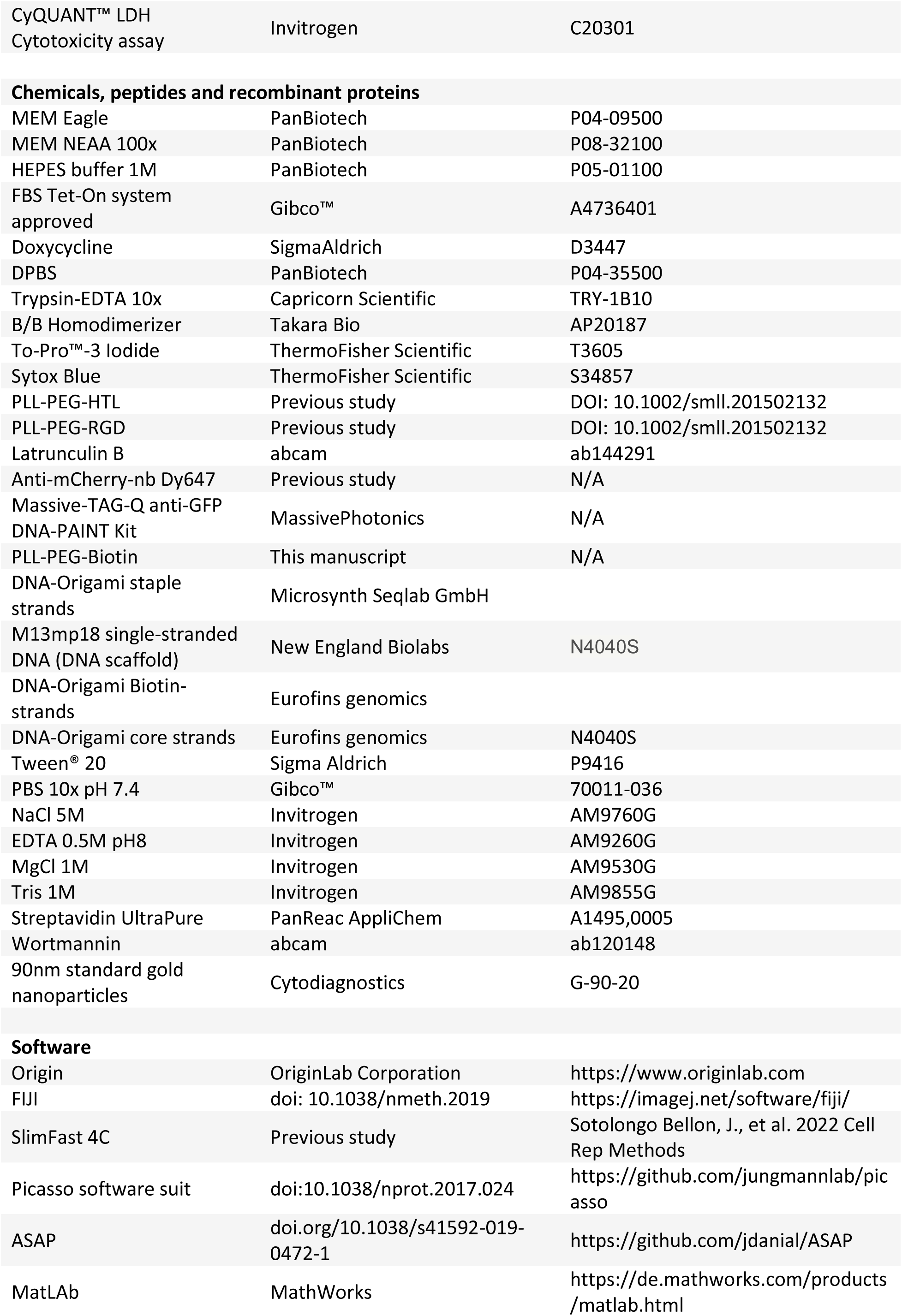
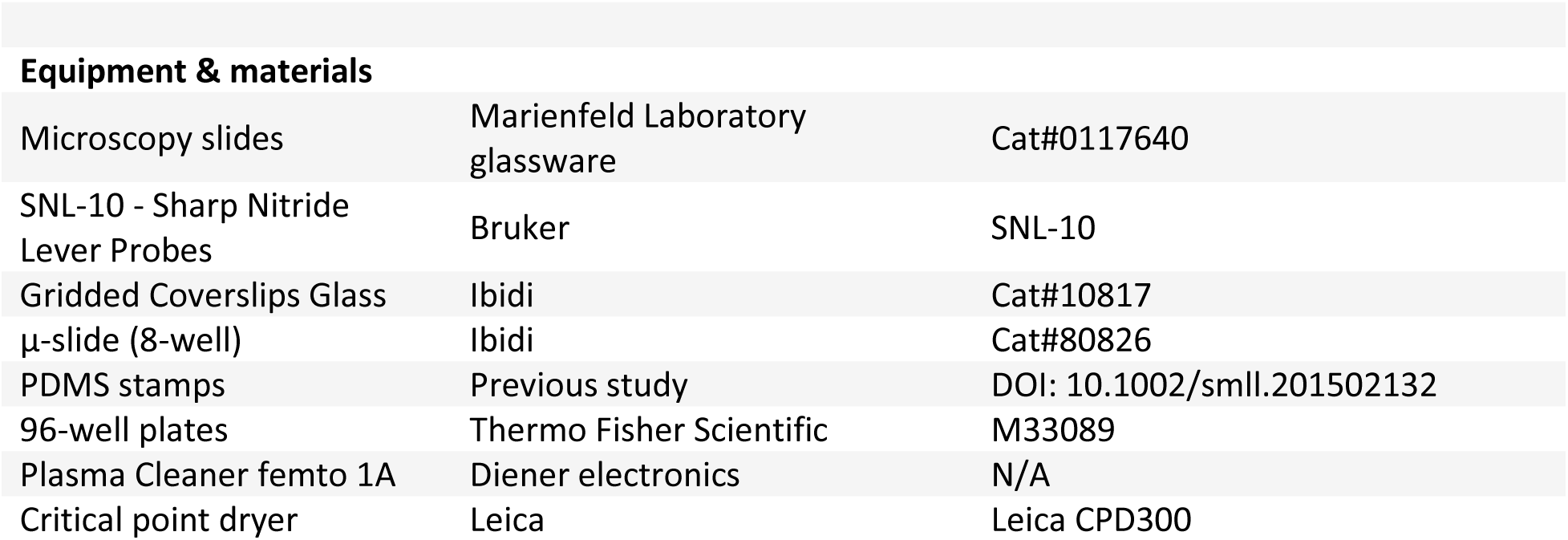

## Author contributions

S.K.: Conceptualization, Methodology, Validation, Formal analysis, Investigation, Writing-Original Draft preparation, Writing-Review&Editing, Visualization; M.H.: Methodology, Software, Formal analysis, data curation, Writing-Review&Editing; S.L.S.: Methodology, Software, Formal analysis, Investigation, Data Curation, Writing-Review&Editing, Visualization; E.G.M.: Conceptualization, Methodology, Validation, Investigation, Writing-Review&Editing; H.V.: Investigation; J.S.D.: Methodology, Software, Data curation, Writing-Review&Editing; S.S.: Methodology, Software, Formal analysis; R.F.: Investigation, Formal analysis; O.E.P.: Methodology, Resources, Supervision, Funding acquisition; R.J.: Methodology, Resources, Supervision, Writing-Editing&Reviewing, Funding acquisition; R.K.: Methodology, Software, Resources, Data curation, Writing-Review&Editing, Funding acquisition; G.H.: Methodology, Resources, Software, Supervision, Writing-Editing&Reviewing, Funding acquisition; J.P.: Conceptualization, Methodology, Software, Resources, Writing-Review&Editing, Supervision, Funding acquisition; K.C.: Conceptualization, Methodology, Validation, Resources, Writing-Original Draft, Writing-Review&Editing, Visualization, Supervision, Project administration, Funding acquisition.

## Acknowledgments

We thank Petr Broz for providing HEK293T DmrB-mCas1 cells. This work was supported by German Research Foundation (DFG, SFB 944, P26 and Z, and SFB 1557, P5 and Z2) to K.C., J.P., R.K. and E.O.P.. S.L.S. and G.H. acknowledge funding through the Collaborative Research Center 1507 “Membrane-associated Protein Assemblies, Machineries, and Supercomplexes“ – Project ID 450648163, thank the Max Planck Society for support, and the Max Planck Computing and Data Facility (MPCDF) for computing resources. S.S and R.J. thank the Max Planck Society for support.

## EXTENDED FIGURES

**Extended Figure 1.**
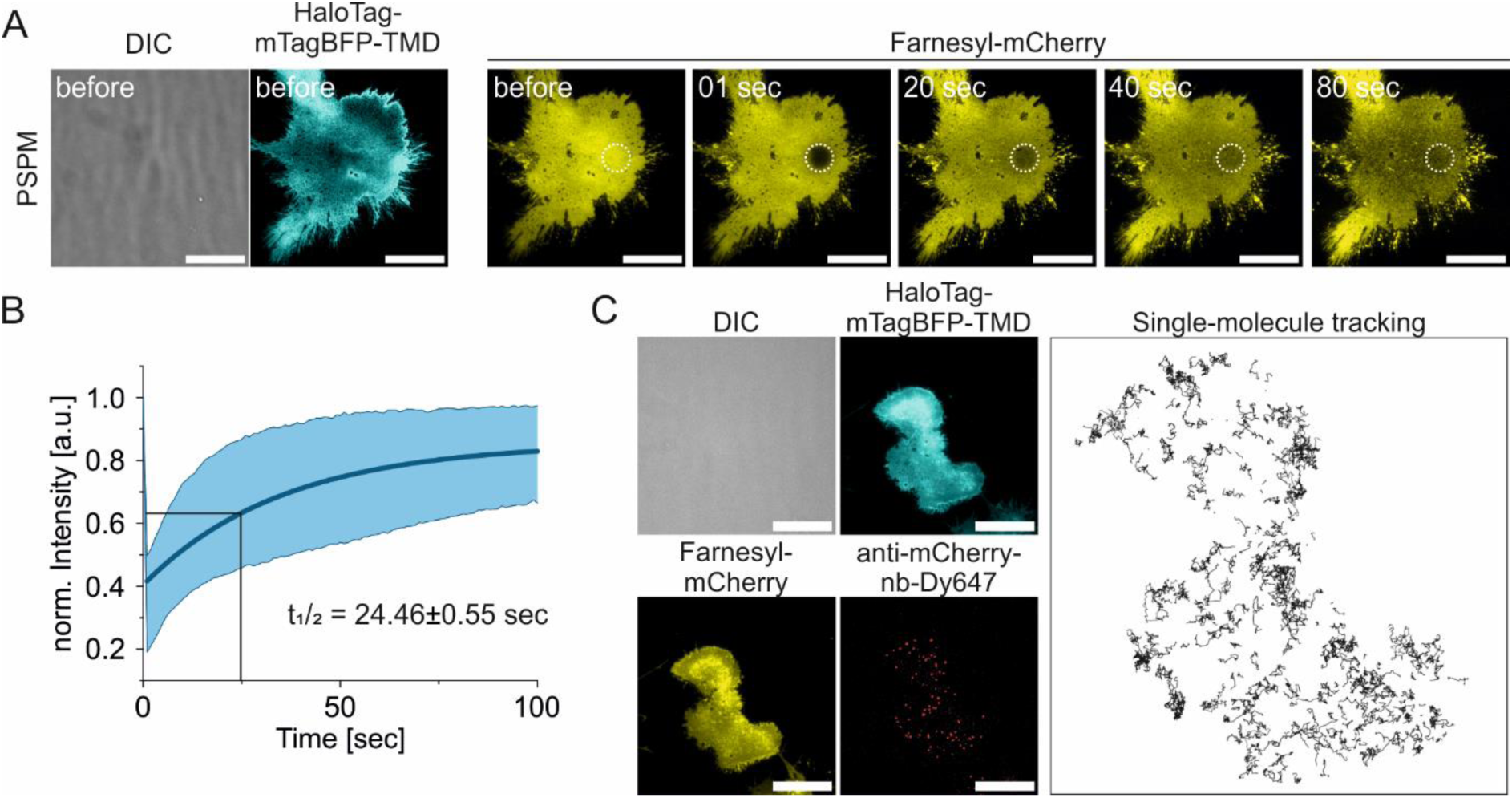
PSPMs maintain membrane fluidity. *(A) FRAP image sequence (TIRFM) before and at 1, 20, 40 and 80 sec after bleaching of the membrane marker Farnesyl-mCherry (yellow) in a PSPM tethered with the HaloTag-mTagBFP-TMD anchor (blue). FRAP region is indicated (white dashed circle). Scale bar 20µm*. *(B) Fluorescence recovery curve of the membrane marker Farnesyl-mCherry in PSPMs (representative images shown in (A)). Mono-exponential fitting of the recovery curve revealed an average recovery half time of 24.5±0.6 s (n=7 cells) resulting in an estimated diffusion constant of 0.23±0.01 µm^2^/s. Line in the graph corresponds to the average value from all measured cells and colored areas to data variability (mean ± SD)*. *(C) Single-molecule tracking of individual anti-mCherry-nb-Dy647-labelled (red) Farnesyl-mCherry proteins (yellow) in PSPMs (blue). The image on the right shows trajectories of individual Farnesyl-mCherry proteins over the whole PSPM. Single-molecule tracking revealed an average diffusion of 0.37±0.05 µm^2^/s. Scale bar 20µm*.

**Extended Figure 2.**
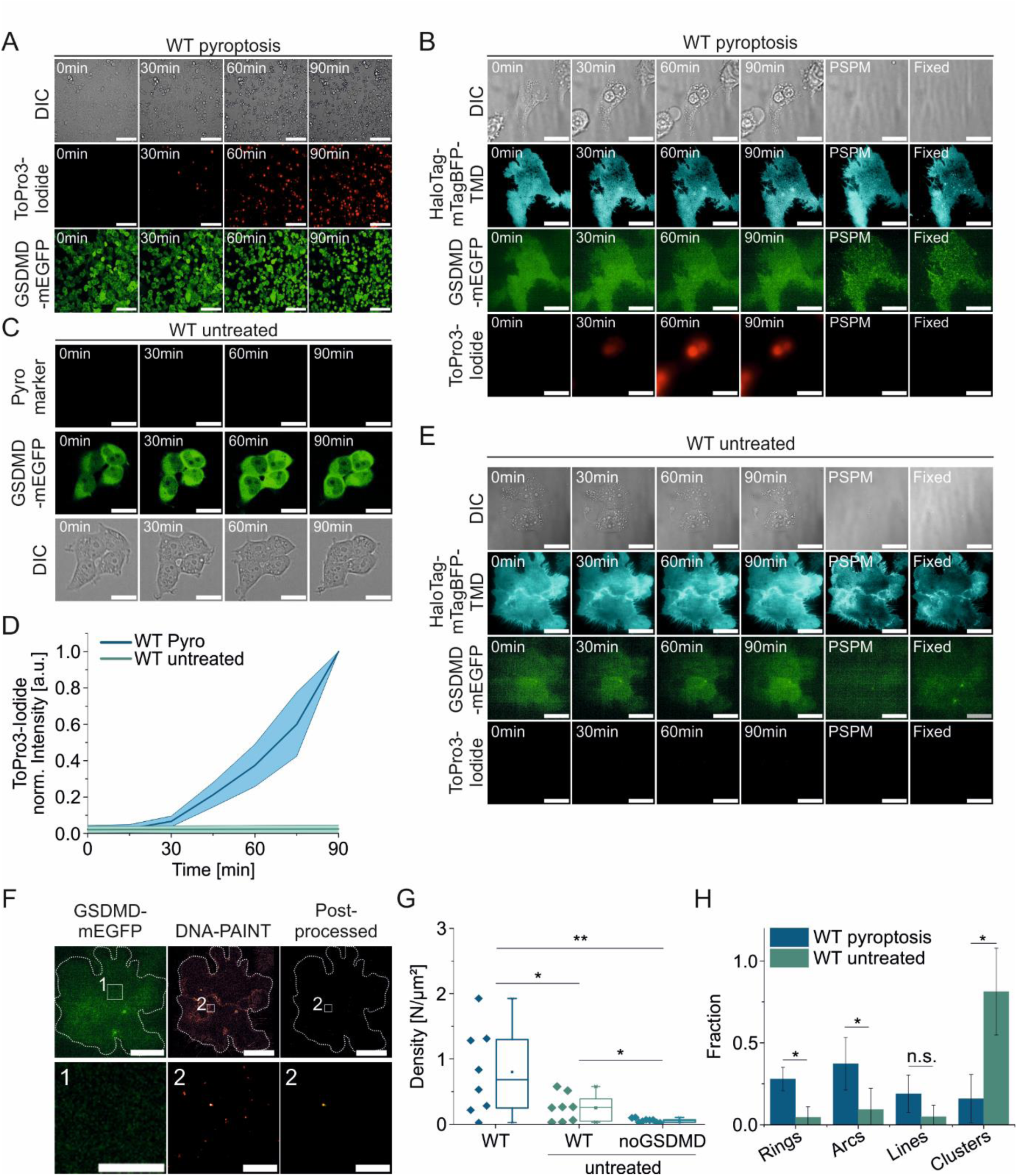
Plasma membrane permeabilization and GSDMD-mEGFP oligomer formation in HEK293T cells upon pyroptosis induction. *(A) Representative confocal fluorescence microscopy images of PM permeabilization (monitored by ToPro3-Iodide uptake, red) after pyroptosis induction at 37°C over 90min in HEK293T cells stably expressing DmrB-mCas1 and mGSDMD-mEGFP (green). Scale bar 100µm* *(B) Representative TIRF microscopy images of a HEK293T cell stably expressing DmrB-mCas1 and mGSDMD-mEGFP upon pyroptosis induction indicated by ToPro3-Iodide staining (red), morphological changes (DIC) and mGSDMD-mEGFP oligomer formation (green), and subsequent PSPM formation (HaloTag-mTagBFP-TMD anchor, blue) and fixation for DNA-PAINT. Scale bar 20µm*. *(C) Representative confocal fluorescence microscopy images of untreated HEK293T cells stably expressing DmrB-mCas1 and mGSDMD-mEGFP WT (green) (PM permeabilization monitored by ToPro3-Iodide uptake, red). Scale bar 20µm*. *(D) Quantification of PM permeabilization of HEK293T cells stably expressing DmrB-mCas1 and mGSDMD-mEGFP WT with and without pyroptosis induction by normalized fluorescence intensity of ToPro3-Iodide (n= 3 experiments with >20 cells analyzed per experiment). Lines in the graph correspond to the average values from all measured cells and colored areas to data variability (mean ± SD)*. *(E) Representative TIRF microscopy images of an untreated HEK293T cell (DIC) stably expressing DmrB-mCas1 and mGSDMD-mEGFP (green) monitored every 30min over 90min with ToPro3-Iodide staining (red) and subsequent PSPM formation (HaloTag-mTagBFP-TMD anchor, blue) and fixation for DNA-PAINT. Scale bar 20µm*. *(F) Upper row: overview of mGSDMD-mEGFP (left) and DNA-PAINT localizations before (middle) and after post-processing (right) of a PSPM from an untreated Hek cmv cell (Scale bars 20µm). Lower row: Zoom in of GSDMD-mEGFP (green) from area 1 in the upper row (left, scale bar 5 µm), zoom in of DNA-PAINT localizations from area 2 in the upper row before (middle) and after (right) post-processing. Scale bar 200 nm*. *(G) Density of super-resolved GSDMD structures after post-processing of DNA-PAINT localizations in HEK293T cells stably expressing either only DmrB-mCas1 or also mGSDMD-mEGFP with or without pyroptosis induction (GSDMD WT with pyroptosis induction n=8 cells; 3 experiments, GSDMD WT untreated n=8 cells; 3 experiments, p=0.041; without GSDMD-mEGFP n=9 cells, 3 experiments, p=0.004(compared to WT treated) and p=0.016(compared to WT untreated))*. *(H) Comparison between the relative distribution of the different GSDMD structure types in HEK293T cells stably expressing DmrB-mCas1 and mGSDMD-mEGFP WT with (n=8 cells, 3 experiments, 493 structures) or without pyroptosis induction (n=8 cells, 3 experiments, 332 structures) (Rings: p=0.039; Arcs: p=0.036; Lines: p=0.227; Undefined clusters: p=0.013)*.

**Extended Figure 3.**
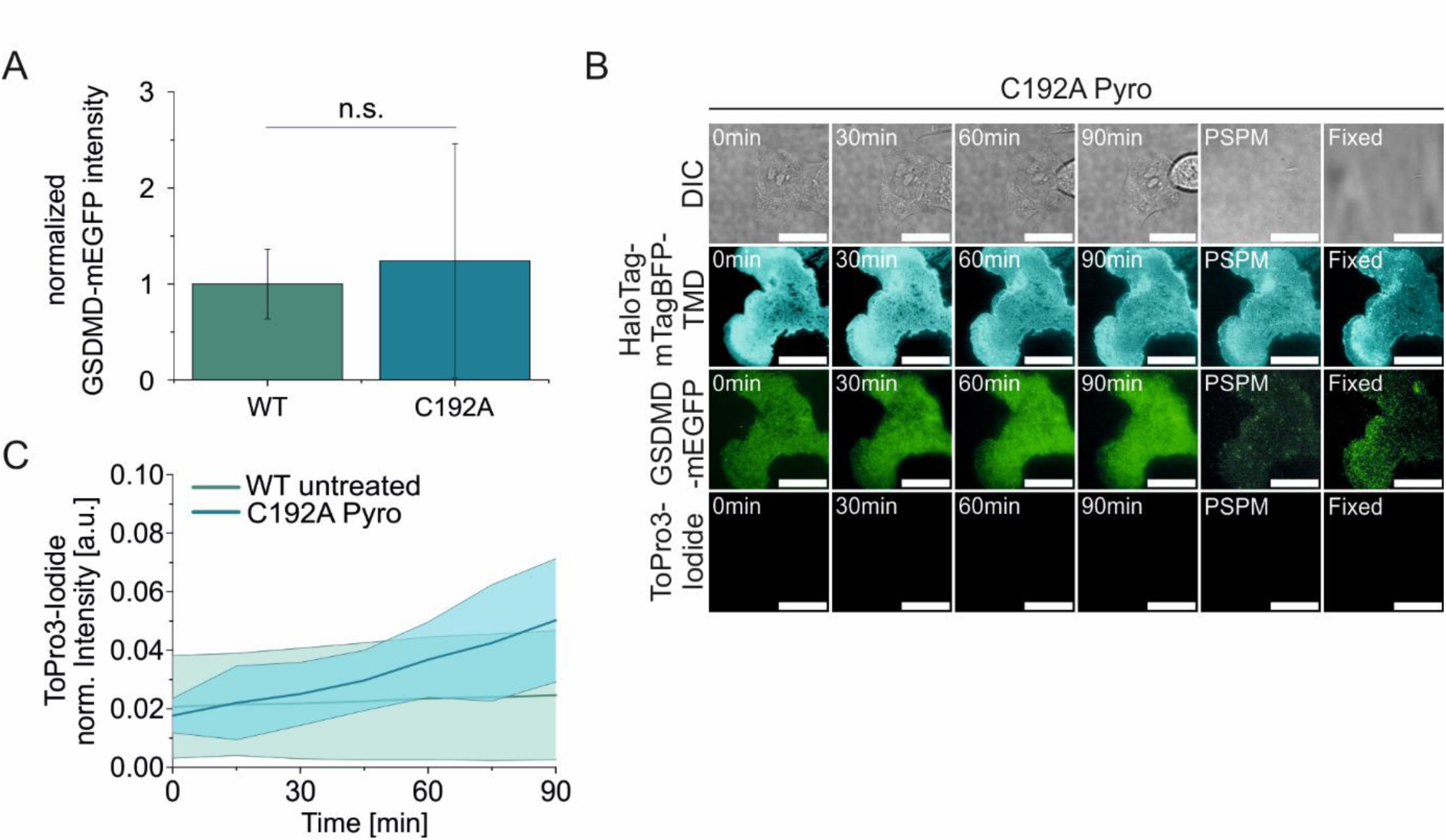
GSDMD-C192A shows PM binding but only slight cell permeabilization. *(A) Difference in the mGSDMD-mEGFP expression level between stable expression of GSDMD-mEGFP WT and transient expression of mGSDMD-C192A-mEGFP measured by the EGFP intensity (p=0.56) (GSDMD-WT: 3 experiments, 238 cells) (GSDMD-C192A: 3 experiments, 105 cells)*. *(B) Representative TIRF microscopy images of a HEK293T cell stably expressing DmrB-mCas1 and mGSDMD-C192A-mEGFP during pyroptosis indicated by ToPro3-Iodide staining (red), morphological changes (DIC) and mGSDMD-C192A-mEGFP oligomer formation (green), and subsequent PSPM formation (HaloTag-mTagBFP-TMD anchor, blue) and fixation for DNA-PAINT. Scale bar 20µm*. *(C) Quantification of PM permeabilization in HEK293T cells stably expressing DmrB-mCas1 and mGSDMD-mEGFP WT without pyroptosis induction or mGSDMD-C192A-mEGFP with pyroptosis induction by normalized fluorescence intensity of ToPro3-Iodide (n= 3 experiments with >20 cells analyzed per experiment). Lines in the graph correspond to the average values from all measured cells and colored areas to data variability (mean ± SD)*.

**Extended Figure 4.**
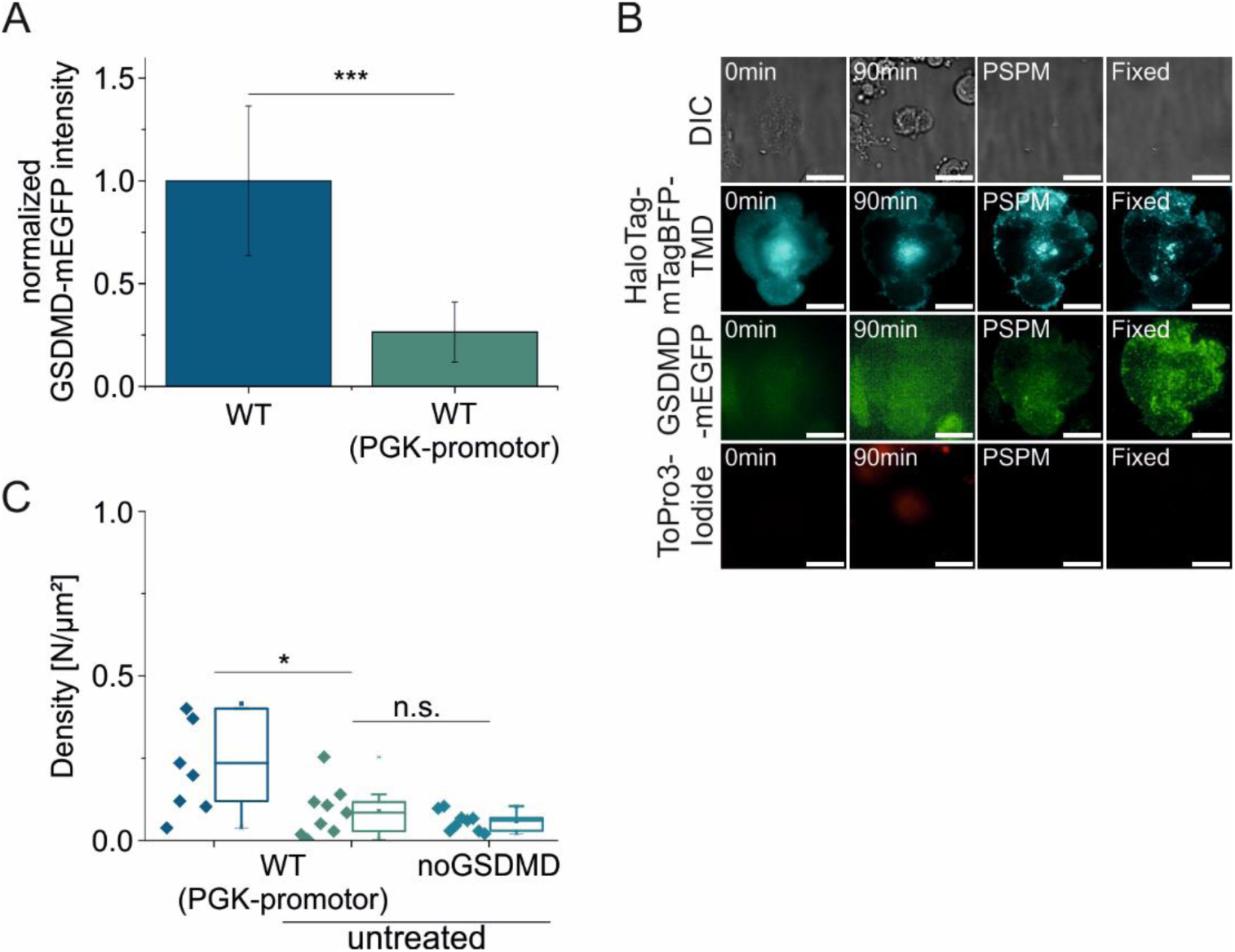
GSDMD at the PM in conditions of low expression of mGSDMD-GFP. *(A) Difference in the GSDMD-mEGFP expression level between a CMV- (WT, used throughout this study) or a PGK-promotor measured by mEGFP intensity (p=0.00007) (CMV: 3 experiments, 238 cells) (PGK: 2 experiments, 170 cells)*. *(B) Representative TIRF images of a HEK293T cell stably expressing DmrB-Cas1 and mGSDMD-mEGFP WT with a PGK-promotor upon pyroptosis induction indicated by ToPro3-Iodide staining (red), morphological changes (DIC) and GSDMD-mEGFP oligomer formation (green), and subsequent PSPM formation (HaloTag-mTagBFP-TMD anchor, blue) and fixation for DNA-PAINT. Scale bar 20µm*. *(C) Density of super-resolved GSDMD structures after post-processing of DNA-PAINT localizations in HEK293T cells stably expressing either only DmrB-mCas1 or also mGSDMD-mEGFP WT (PGK-promotor) with and without pyroptosis induction (GSDMD WT (PGK-promotor) with pyroptosis induction n=7 cells; 3 experiments, GSDMD WT (PGK-promotor) untreated n=9 cells; 3 experiments, p=0.038; without GSDMD-mEGFP n=9 cells, 3 experiments, p=0.28(comp.to WT untreated))*.

**Extended Figure 5.**
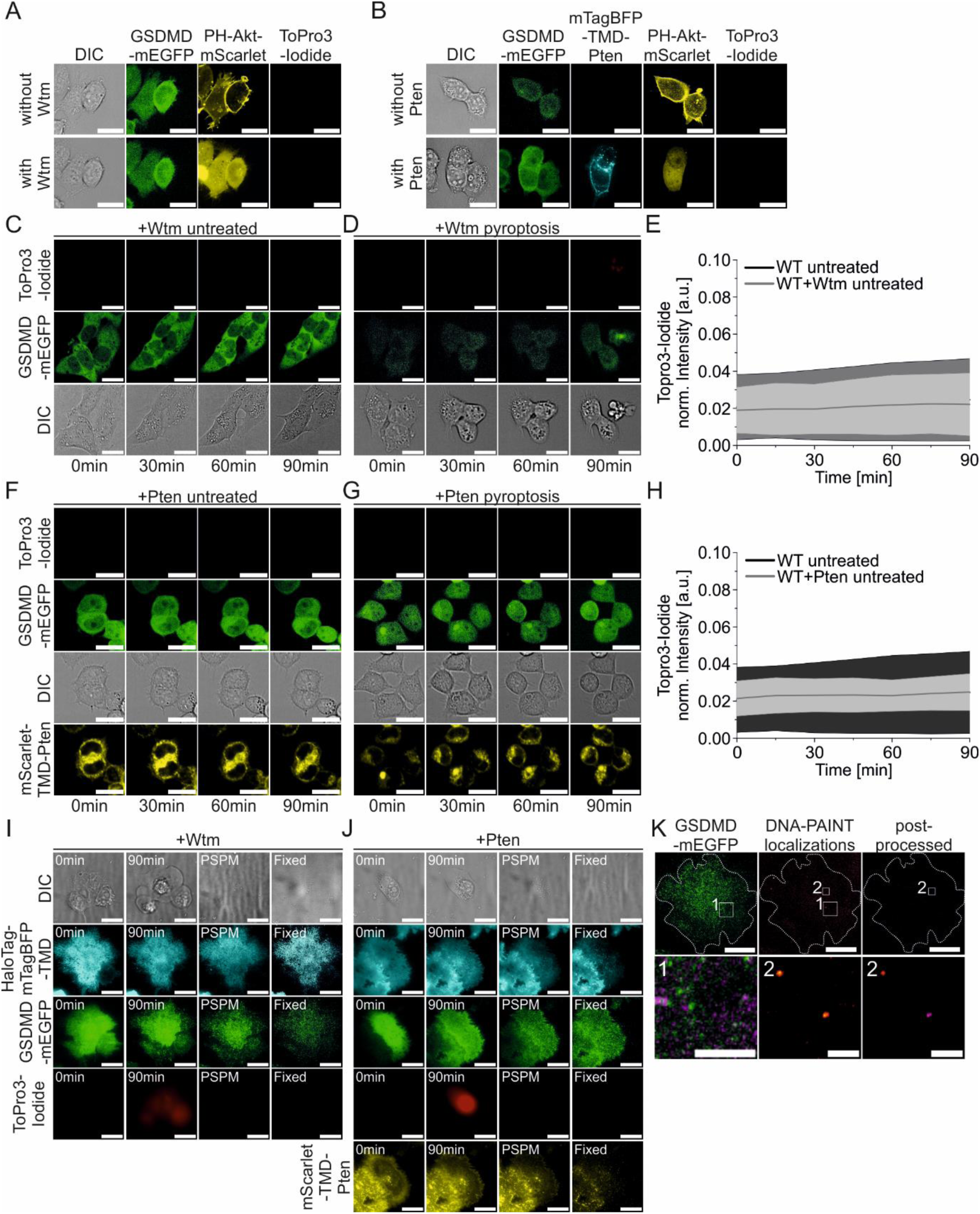
PI(3,4,5)P_3_ depletion by Wtm treatment or Pten expression alone has no consequence on cell permeabilization. *(A) Representative confocal images of PI(3,4,5)P_3_ levels (PH-Akt-mScarlet, yellow) at the PM of non-pyroptotic Hek293T cells (PM permeabilization monitored by ToPro3-Iodide uptake, red) stably expressing DmrB-mCas1 and mGSDMD-mEGFP (green) before (upper row) and after (lower row) treatment with 10µM Wtm for 10min. Scale bar 20µm*. *(B) Representative confocal images of PI(3,4,5)P_3_ levels (PH-Akt-mScarlet, yellow) at the PM of non-pyroptotic Hek293T cells (PM permeabilization monitored by ToPro3-Iodide uptake, red) stably expressing DmrB-mCas1 and mGSDMD-mEGFP (green) without (upper row) or with (lower row) expression of mTagBFP-TMD-Pten (cyan). Scale bar 20µm*. *(C) Representative confocal fluorescence microscopy images of HEK293T cells stably expressing DmrB-mCas1 and mGSDMD-mEGFP WT (green) without pyroptosis induction but with treatment of 10µM Wtm (PM permeabilization monitored by ToPro3-Iodide uptake, red). Scale bar 20µm*. *(D) Representative confocal fluorescence microscopy images of HEK293T cells stably expressing DmrB-mCas1 and mGSDMD-mEGFP WT (green) with pyroptosis induction and treatment with 10µM Wtm (PM permeabilization monitored by ToPro3-Iodide uptake, red). Scale bar 20µm*. *(E) Quantification of PM permeabilization in HEK293T cells stably expressing DmrB-mCas1 and mGSDMD-mEGFP WT without pyroptosis induction and with or without 10µM Wtm treatment by normalized fluorescence intensity of ToPro3-Iodide (n= 3 experiments with >20 cells analyzed per experiment). Lines in the graph correspond to the average values from all measured cells and colored areas to data variability (mean ± SD)*. *(F) Representative confocal fluorescence microscopy images of HEK293T cells stably expressing DmrB-mCas1 and mGSDMD-mEGFP WT (green) and transiently expressing mScarlet-TMD-Pten (yellow) without pyroptosis induction (PM permeabilization monitored by ToPro3-Iodide uptake, red). Scale bar 20µm*. *(G) Representative confocal fluorescence microscopy images of HEK293T cells stably expressing DmrB-mCas1 and mGSDMD-mEGFP WT (green) and transiently expressing mScarlet-TMD-Pten (yellow) upon pyroptosis induction (PM permeabilization monitored by ToPro3-Iodide uptake, red). Scale bar 20µm*. *(H) Quantification of PM permeabilization in HEK293T cells stably expressing DmrB-mCas1, mGSDMD-mEGFP WT and transiently expressing mScarlet-TMD-Pten without pyroptosis induction by normalized fluorescence intensity of ToPro3-Iodide over 90min (n= 3 experiments with >20 cells analyzed per experiment). Lines in the graph correspond to the average values from all measured cells and colored areas to data variability (mean ± SD)*. *(I) Representative TIRF images of a HEK293T cell stably expressing DmrB-Cas1 and mGSDMD-mEGFP WT treated with 10µM Wtm upon pyroptosis induction indicated by ToPro3-Iodide staining (red), morphological changes (DIC) and GSDMD-mEGFP oligomer formation (green), and subsequent PSPM formation (HaloTag-mTagBFP-TMD anchor, blue) and fixation for DNA-PAINT. Scale bar 20µm*. *(J) Representative TIRF images of a HEK293T cell stably expressing DmrB-Cas1, GSDMD-mEGFP WT and transiently expressing mScarlet-TMD-Pten (yellow) upon pyroptosis induction indicated by ToPro3-Iodide staining (red), morphological changes (DIC) and GSDMD-mEGFP oligomer formation (green), and subsequent PSPM formation (HaloTag-mTagBFP-TMD anchor, blue) and fixation for DNA-PAINT. Scale bar 20µm*. *(K) Upper row: overview of GSDMD-mEGFP and DNA-PAINT localizations before and after post-processing of a PSPM from a pyroptotic cell treated with 10µm Wtm (Scale bars 20µm). Lower row: Correlation of GSDMD-mEGFP (green) and DNA-PAINT localizations (magenta) from zoom-in area 1 in the upper row (left, scale bar 5 µm), zoom in of DNA-PAINT localizations from area 2 in the upper row before (middle) and after (right) post-processing. Scale bar 200 nm*.

**Extended Figure 6.**
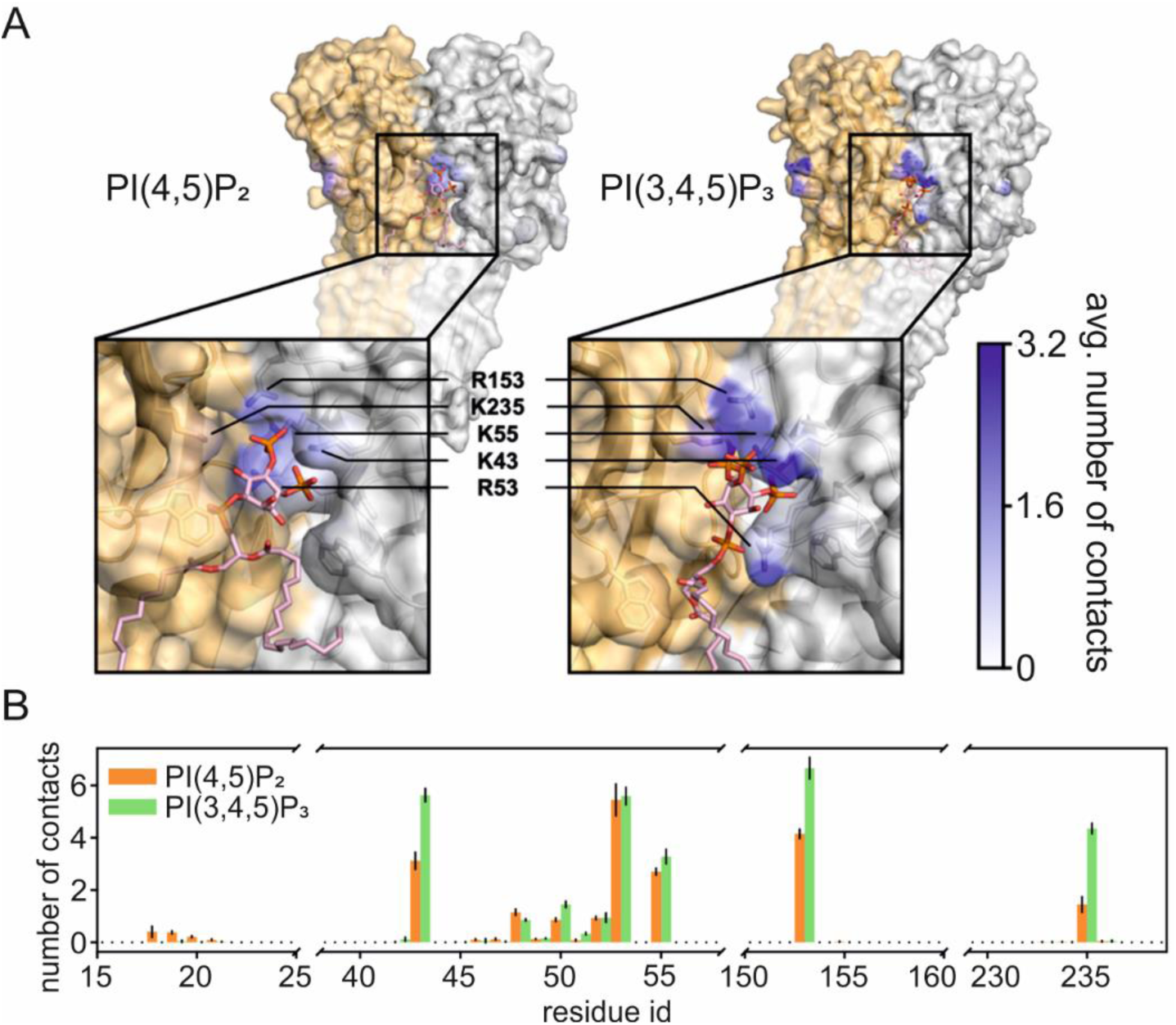
PI(3,4,5)P3 binds to 5 positively charged residues in the interface of GSDMD dimers. *(A) Snapshots of human GSDMD dimers extracted from the final frame of the 16-mer simulations with PI(4,5)P2 or PI(3,4,5)P3. The protein is shown in white surface representation with PI(4,5)P2 (left) and PI(3,4,5)P3 (right) binding sites shown and colored according to the average number of contacts between the respective PIP species and any protein heavy atoms in the MD simulations. Protein residues with at least one atom that has more than 0.2 average contacts are shown in licorice representation*. *(B) Average number of contacts between PI(4,5)P2 and PI(3,4,5)P3 heavy atoms with each GSDMD residue. Error bars depict the standard deviation of the three replicate simulations. Areas excluded by the cut x-axis did not exhibit any contacts with either PIP species*.

*Attached media:*

***Extended Video 1*** *Simulation of a 16-mer arc-shaped GSDMD oligomer in a pure DOPC membrane*.

*Simulation over the time course of 600 ns. GSDMD subunits (blue) and DOPC lipids (yellow). Black triangles indicate the cracks between GSDMD subunits created by the membrane tension*.

***Extended Video 2*** *Simulation of a 16-mer arc-shaped GSDMD oligomer in a DOPC membrane with PI(4,5)P2*.

*Simulation over the time course of 600 ns. GSDMD subunits (blue), DOPC lipids (yellow) and PI(4,5)P2 (orange). Black triangles indicate the cracks between GSDMD subunits created by the membrane tension*.

***Extended Video 3*** *Simulation of a 16-mer arc-shaped GSDMD oligomer in a DOPC membrane with PI(3,4,5)P3.*

*Simulation over the time course of 600 ns. GSDMD subunits (blue), DOPC lipids (yellow) and PI(3,4,5)P3 (green). Black triangles indicate the cracks between GSDMD subunits created by the membrane tension*

